# Oriented basement membrane fibrils provide a memory for F-actin planar polarization via the Dystrophin-Dystroglycan complex during tissue elongation

**DOI:** 10.1101/591628

**Authors:** Fabiana Cerqueira Campos, Cynthia Dennis, Hervé Alégot, Cornelia Fritsch, Adam Isabella, Pierre Pouchin, Olivier Bardot, Sally Horne-Badovinac, Vincent Mirouse

## Abstract

How extracellular matrix participates to tissue morphogenesis is still an open question. In the *Drosophila* ovarian follicle, it has been proposed that after Fat2-dependent planar polarization of the follicle cell basal domain, oriented basement membrane (BM) fibrils and F-actin stress fibers constrain follicle growth, promoting its axial elongation. However, the relationship between BM fibrils and stress fibers and their respective impact on elongation are unclear. We found that Dystroglycan (Dg) and Dystrophin (Dys) are involved in BM fibril deposition. Moreover, they orient stress fibers, by acting locally and in parallel to Fat2. Nonetheless, Dg-Dys complex-mediated cell autonomous control of F-actin fibers orientation relies on the previous BM fibril deposition, indicating two distinct but interdependent functions. Thus, the Dg-Dys complex works as a critical organizer of the epithelial basal domain, regulating both F-actin and BM. Furthermore, BM fibrils act as a persistent cue for the orientation of stress fibers that are the main effector of elongation.

## Introduction

Deciphering the mechanisms underlying tissue morphogenesis is crucial for our fundamental understanding of development and also for regenerative medicine. Building organs generally requires the precise modeling of a basement membrane extracellular matrix (ECM), which in turn can influence tissue shape (Dzamba and DeSimone, 2018; Sekiguchi and Yamada, 2018; Isabella and Horne-Badovinac, 2015). However, the mechanisms driving the assembly of a specific basement membrane (BM) and how this BM then feeds forward on morphogenesis are still poorly understood. *Drosophila* oogenesis offers one of the best tractable examples in which such a morphogenetic process. Each ovarian follicle, which is composed of a germline cyst surrounded by the somatic follicular epithelium undergoes a dramatic growth, associated with tissue elongation, starting from a little sphere and ending with an egg in which the anteroposterior (AP) axis is 3-fold longer than the mediolateral (ML) axis (Fig. 1A). This elongation is roughly linear from the early to the late stages, but can be separated in at least two mechanistically distinct phases (Alégot et al., 2018; Aurich and Dahmann, 2016). The first phase (from stage 3 to stage 8, hereby “early stages”) requires a double gradient of JAK-STAT pathway activity that emanates from each pole and that controls myosin II-dependent apical pulsations (Alégot et al., 2018). In the second phase, from stage 7-8, elongation depends on the atypical cadherin Fat2 that is part of a planar cell polarity (pcp) pathway orienting the basal domain of epithelial follicle cells (Gutzeit et al., 1991; Barlan et al., 2017; Chen et al., 2016; Viktorinová et al., 2009). Earlier during oogenesis, Fat2 gives a chirality to the basal domain cytoskeleton in the germarium, the structure from which new follicles bud (Chen et al., 2016). This chirality is required to set up a process of oriented collective cell migration perpendicularly to the elongation axis that induces follicle revolutions from stage 1 to stage 8 (Chen et al., 2016; Viktorinová and Dahmann, 2013). From each migrating cell, Fat2 also induces, in the rear adjacent cell, the formation of planar polarized protrusions that are required for rotation (Cetera et al., 2014, Barlan et al., 2017). These rotations allow the polarized deposition of BM fibrils, via a secretion route to the lateral domain of the cells, implicating Rab10. These BM fibrils are detectable since stage 4 and persist until late developmental stages (Haigo and Bilder, 2011; Isabella and Horne-Badovinac, 2016). Follicle rotation also participates to the planar cell polarization of integrin-dependent basal stress fibers that are oriented perpendicularly to the AP axis (Cetera et al., 2014). Moreover, at stage 7-8, a gradient of matrix stiffness controlled by the JAK-STAT pathway and Fat2 contributes to elongation (Crest et al., 2017). Then, from stage 9, the epithelial cell basal domain undergoes anisotropic oscillations, due to the periodic contraction of the oriented stress fibers, which also promotes follicle elongation (He et al., 2010; Qin et al., 2017). To explain the impact of *fat2* mutations on tissue elongation, it is generally accepted that oriented stress fibers and BM fibrils act as a molecular corset that constrains follicle growth in the ML axis and promotes its elongation along the AP axis. However, the exact contribution of F-actin versus BM to this corset is still unclear, as is whether the orientations of stress fibers and of BM fibrils are causally linked.

**Figure 1:**
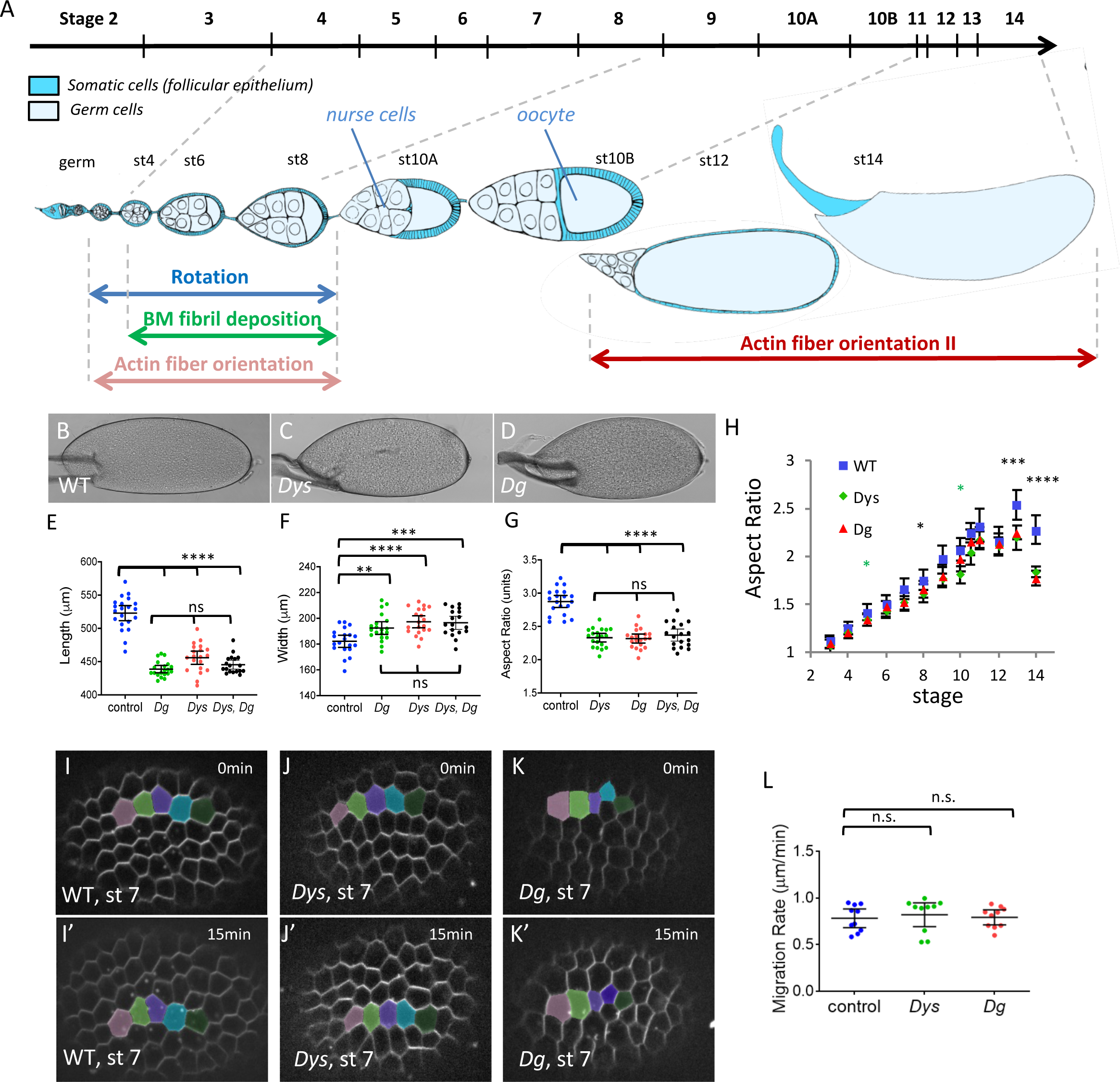
The DAPC is involved in follicle elongation but not rotation. A) Scheme of an ovariole with the main events involved in follicle elongation. The above line indicates the time scale of the different developmental stages. The ovariole is oriented anterior to posterior (germ: germarium). Each follicle is composed of a germline cyst surrounded by the follicular epithelium. Rotation occurs from very early to stage 8. It promotes F-actin fiber orientation and allows polarized BM fibril deposition, both perpendicularly to the elongation axis. At stage 11, actin fibers lose their orientation and then progressively reorient (orientation II). B,C,D) Representative mature eggs from B) WT, C) *Dys^E17^*^/*Exel6184*^ and D) *Dg^O46/O83^* females. E,F,G) Quantification of E) the length, F) the width, and G) the aspect-ratio for mature eggs from WT, *Dys^E17^*^/ *Exel6184*^, *Dg^O46/O83^* mutant and *Dys^E17^*^/ *Exel6184*^; *Dg^O46/O83^* double mutant females (n>18 eggs). H) Elongation kinetics of WT, *Dys ^E17^*^/*Exel6184*^ and *Dg ^O46/O83^* mutant follicles (n>6 follicles for each point, green stars for *Dys*, Black stars for *Dys* and *Dg).* There are two series of value for stage 10, corresponding to stage 10A and 10B. I,J,K) Images of rotation movies of stage 7 I) WT, J) *Dys^E17^*^/ *Exel6184*^ and K) *Dg^O46/O83^* mutant follicles. L) Rotation velocity of WT, *Dys ^E17^*^/*Exel6184*^ and *Dg ^O46/O83^* mutant stage 7 follicles (n>8 follicles). (For all panels error bars represent s.d.; p *< 0.01, **<0.005, ***<0.001, ****<0.0001)

Here, we analyzed the function of Dystrophin (Dys) and Dystroglycan (Dg) during follicle elongation. Dys and Dg are the two main components of the Dystrophin-Associated Protein Complex (DAPC), an evolutionarily conserved transmembrane complex that links the ECM (via Dg) to the F-actin cytoskeleton (via Dys) (Barresi and Campbell, 2006). This complex is expressed in a large variety of tissues and is implicated in many congenital degenerative disorders. Loss of function studies in model organisms have revealed an important morphogenetic role for Dg during development, usually linked to defects in ECM secretion, assembly or remodeling (Clements et al., 2017; Henry and Campbell, 1998; Satz et al., 2010; Bello et al., 2008; Buisson et al., 2014; Naegeli et al., 2017; Yatsenko and Shcherbata, 2014). A developmental role for Dys is less clear, possibly due to the existence of several paralogs in vertebrates. As *Drosophila* has only one *Dg* and one *Dys* gene, it is a promising model for their functional study during development and morphogenesis.

We found that *Dg* and *Dys* are required for follicle elongation and proper BM fibril formation early in fly oogenesis. During these early stages, DAPC loss and hypomorphic *fat2* conditions similarly delay stress fiber orientation. However, DAPC promotes this alignment more locally than Fat2. Moreover, DAPC genetically interacts with *fat2* in different tissues, suggesting that they belong to a common morphogenetic network. Later in oogenesis, Dg and Dys are required for stress fiber orientation in a cell autonomous manner. This is the period when the main elongation defect is seen in these mutants, arguing for a predominant role for stress fibers in the elongation process. Nonetheless, this latter function depends on the earlier DAPC function in BM fibril deposition. We propose that BM fibrils serve as a pcp memory for the late stages, and are used as a template by the DAPC for F-actin stress fiber alignment.

## Results

### *Dys* and *Dg* mutants reveal a new phase of follicle elongation

As in *Drosophila* follicle elongation requires many proteins that function at the basal side of the follicular epithelium, we checked whether the DAPC could be involved in this process. Both *Drosophila Dg* and *Dys* null mutants are viable and we used combinations of two different null alleles for each of them (Christoforou et al., 2008). Mature eggs from mutant females are significantly shorter and wider (Fig. 1B-F) with an aspect ratio (AR) around 2.3 compared with 2.9 in wild type (WT) follicles (Fig. 1G). Importantly, such a phenotype can be rescued by restoring expression in the follicle cells, confirming that it is associated with these mutations, and indicating that the DAPC is required for elongation in the follicle cells (see below). Moreover, double mutant eggs for *Dy*s and *Dg* showshowed the same defect than single ones confirming that these genes work together for follicle elongation (Fig. 1E-G). Comparison of the elongation kinetics in mutant and WT follicles shows that such morphological differences appear mainly during the later stages of oogenesis (stages 12 to 14), although a slight, but significant defect is present around stage 8 (Fig. 1H). The *Dg* and *Dys* null mutants elongation kinetic curves were clearly different from those of *fat2* mutant in which elongation is strongly affected from stages 7-8 (Alégot et al., 2018; Aurich and Dahmann, 2016). Thus, the DAPC is involved in follicle morphogenesis, and its specific temporal requirement suggests the existence of an elongation phase that is mechanistically different from the Fat2-dependent one.

Then, to determine how the DAPC contributes to follicle elongation, we first checked the collective cell migration that occurs during the early stages because it is upstream of many subsequent events that are potentially important for elongation. Both *Dys* and *Dg* null mutant follicles rotate and the migration velocity is not significantly different from that of WT follicles (Fig. 1I-L). Thus, the DAPC is not involved in the collective cell migration and in the initial pcp established by Fat2, confirming that it acts through a distinct mechanism. We then focused our phenotypic analysis on the two events induced by follicle rotation : BM fibril deposition and stress fiber orientation.

### The DAPC is important for BM fibril deposition

The follicle cell BM contains two ECM types, a planar BM without a specific organization visible by confocal microscopy, and BM fibrils that are oriented perpendicular to the follicle AP axis. BM fibrils are deposited during follicle rotation via a secretory route targeted to the cell lateral domain, and can be observed by monitoring the expression of the main ECM components: Collagen IV (ColIV), Perlecan (Pcan) and Laminin A (LanA), the two latter being direct Dg ligands (Gutzeit et al., 1991; Haigo and Bilder, 2011; Isabella and Horne-Badovinac, 2016). ColIV and Pcan can be visualized using the Viking- and Trol-GFP protein traps, respectively (Morin et al., 2001). Moreover, a fosmid containing the whole *laminin A* gene tagged with GFP has also been generated (LanA-GFP) (Sarov et al., 2016). We also employed the MiMIC system to generate RFP-tagged Pcan (Venken et al., 2011). This line gives an identical pattern to Pcan-GFP and is homozygous viable and fertile, indicating that it does not disrupt Pcan function (not shown). Comparison of the BM fibril composition using these different tagged proteins (Fig. 2A-B) showed that in WT follicles, Pcan-RFP co-localizes exactly with ColIV-GFP and LanA-GFP. This indicates that BM fibrils are uniformly composed of these three proteins, as previously shown by immunostaining (Isabella and Horne-Badovinac, 2016).

**Figure 2:**
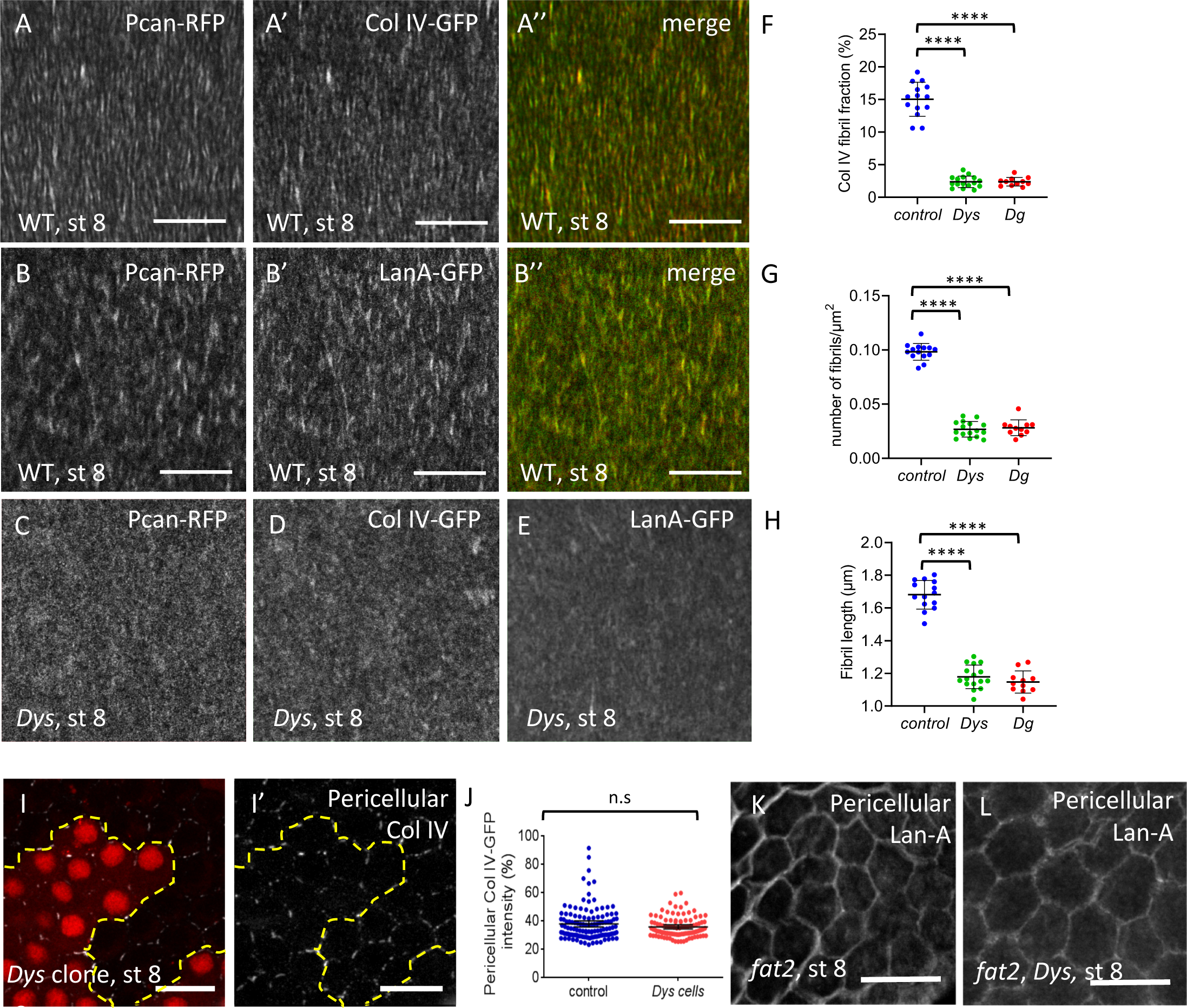
The DAPC is important for BM fibril deposition. A-B) Basal view of WT (*w^1118^*) follicle ECM at stage 8 visualized with A) Pcan-RFP andColIV-GFP and B) Pcan-RFP and LanA-GFP C) Basal view of *Dys^E17/Exel8164^* follicles expressing C) Pcan-RFP D) ColIV-GFP E) LanA-GFP. F-H) Quantification of F) the BM fibril fraction, G) fibril numbers and H) fibril length at stage 8 in WT, *Dys^E17/Exel8164^* and *Dg^O86/O43^* mutant follicles (n> 10 follicles). I) Mutant clones for *Dys^Exel8164^* marked by the absence of RFP and stained to detect pericellular ColIV J) Quantification of pericellular ColIV intensity around WT and *Dys* mutant cells (n> 20 cells). K,L) Accumulation of pericellular LanA in K) *fat2* and L) *fat2, Dys* mutant follicles. For all panels error bars represent s.d.; p *< 0.01, **<0.005, ***<0.001, ****<0.0001. Scalebar 10µm

We then wondered whether BM fibril deposition is affected in *Dys* and *Dg* mutants. In these two mutants, the general targeting of BM secretion towards the basal domain is normal, with no visible apical secretion as it has been observed in different mutants affecting BM secretion (Fig. S2A-C) (Denef et al., 2008; Devergne et al., 2014; Lerner et al., 2013; Devergne et al., 2017). However, images in the focal plane of the BM reveal a very strong decrease of the ColIV-fibrils in *Dys* null mutant (Fig. 2D). This observation is also true for LanA and Pcan (Fig. 2C and E). Quantification of the BM fibril fraction on stage 8 follicles, which corresponds to fluorescence fraction present in the fibrils, indicates that it corresponds to about 15% in the WT situation, but drops to 2% in *Dys* mutants, and that a similar phenotype is observed in *Dg* mutants (Fig. 2F). Loss of the DAPC affects both BM fibril number and length (Fig. 2G,H). As a result of this defect, some fibrils are still visible in stage 12 WT follicles but hardly detectable in *Dys* and *Dg* mutant ones (Fig. S2D-G). This shows that the DAPC is involved in BM fibril deposition.

This deposition depends on the lateral secretion of ECM proteins while the rotation facilitates both the deposition and orientation of the fibrils in the BM (Isabella and Horne-Badovinac, 2016). The lateral secretion of BM proteins can be visualized by immunostaining on non-permeabilized follicle cells to detect only the proteins localized between the cells. Such an approach on *Dys* mutant clones shows that some ColIV is detected, as between WT cells (Fig. 2 I,J). This suggests that Dys does not directly interfere with the lateral targeting of BM fibril secretion. In a mutant blocking rotation, such as *fat2*, BM proteins accumulate in the lateral space between cells, whereas no massive accumulation is observed in a *Dys* mutant, suggesting that the BM fibril defect associated with DAPC loss of function cannot be explained by a failure of the latest step of BM fibril deposition (Fig. 2K) (Isabella and Horne-Badovinac, 2016). Moreover, in a *fat2, Dys* double mutant we observed the same lateral accumulation of ECM proteins as in a *fat2* mutant, confirming that lateral secretion is not affected by DAPC loss of function (Fig. 2 K, L). It is therefore unclear by which mechanism the DAPC contributes to fibril formation. Nonetheless, these mutants are the first examples of mutations that specifically reduce BM fibril formation without any defect in the general targeting of basal secretion or in rotation, offering a unique opportunity to address the functional relevance of the fibrils.

### The DAPC contributes to stress fiber orientation during early stages

In parallel to the analysis of BM fibrils, we also studied the impact of the loss of the DAPC on basal F-actin organization. This alignment is still weak at stage 4 and then increases to reach a maximum around stage 8 (Fig. 3A-F) (Cetera et al., 2014). Both in *Dys* and *Dg* null mutants, although the stress fibers are globally oriented perpendicular to the AP axis in mutant follicles, quantification of the angular distribution reveals a delay in their proper orientation at stage 4 and 6 to reach a normal level at stage 8 (Fig. 3A-R). Stress fiber orientation can be also analysed by the determination of the Tissue Order Parameter (Tissue OP) based on a method previously described (Cetera et al., 2014). A complete random orientation Tissue OP value will tend to 0, whereas a perfect alignment between the cells will tend to 1 (Fig. 3S). This method confirms that *Dys* and *Dg* mutants behave similarly, with a delay in the proper stress fiber alignment (Fig. 3T). Although rotation is required for stress fiber orientation, this motion is not affected by DAPC loss of function. Thus, this complex likely acts downstream of or in parallel to rotation to orient stress fibers.

**Figure 3:**
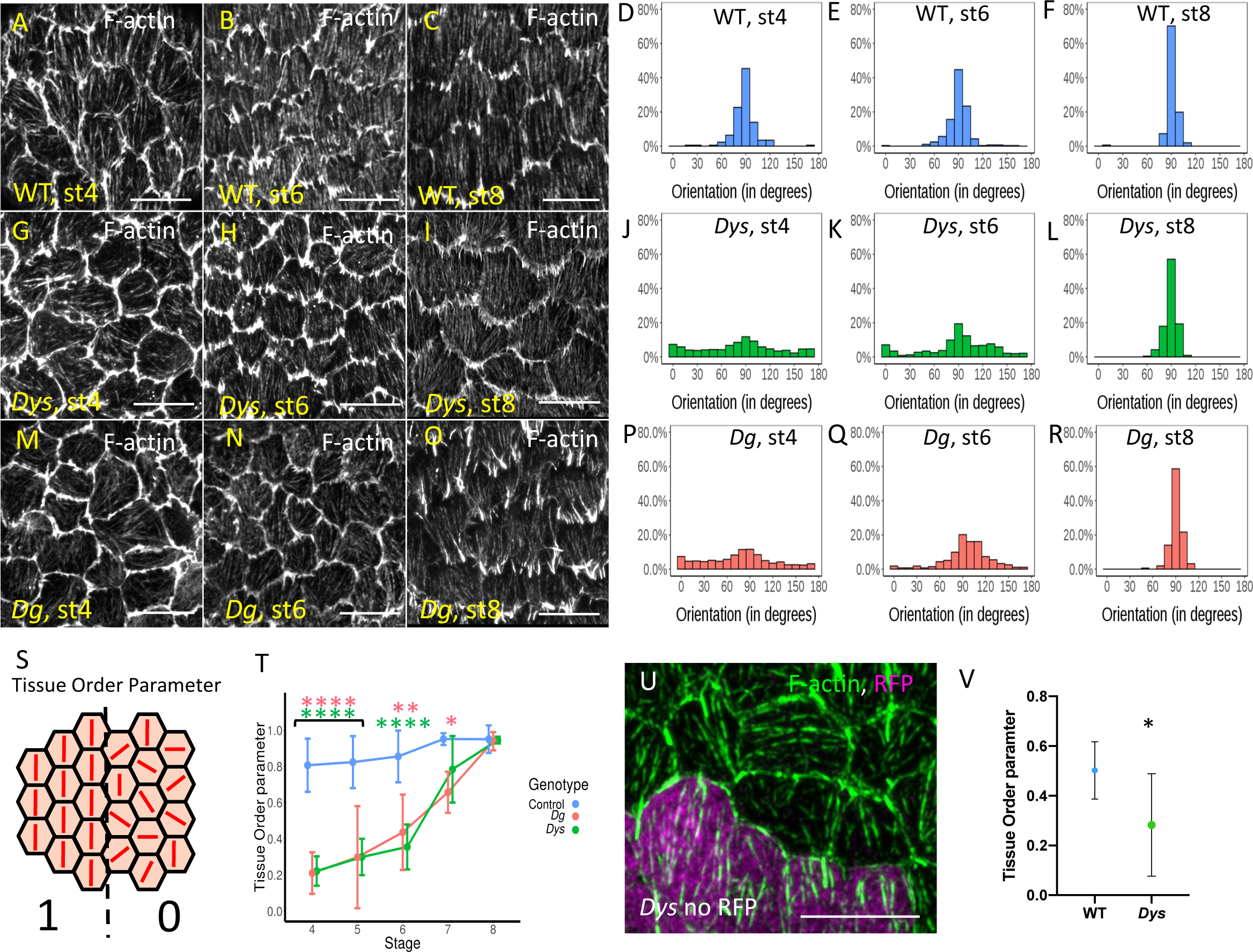
Stress fiber orientation delay in DAPC mutants during early stages. Representative images of basal F-actin in A,B,C) WT (*w^1118^*), G,H,I) *Dys^E17/Exel8164^* and M,N,O) *Dg^O86^/^O43^* follicles at stage 4, 6 and 8 and D,E,F,J,K,L,P,Q,R) Quantification of the corresponding angular distribution. S) Illustration of Tissue Order Parameter (Tissue OP) : if all the cells have the same orientation Tissue OP is equal to one, if orientation is random Tissue OP is equal to zero. T) Tissue OP analysis of control, *Dys ^E17^*^/*Exel6184*^ and *Dg ^O46/O83^* follicles. U) F-actin in a mutant clone for Dys^Exel6184^ marked by the absence of RFP (purple) at stage 5. Note the cell autonomous effect on stress fiber orientation. V) Tissue OP quantification Dys^Exel6184^ mutant clones and surrounding WT cells at stage 5. Error bars represent s.d (n> 7 follicles for each stage and each genotype). p *< 0.01, **<0.005, ***<0.001, ****<0.0001. Scalebar 10µm

We also addressed whether *Dys* is required cell autonomously to promote stress fiber orientation by generating mutant clones. We focused on stage 5, when the defect is obvious in *Dys* mutant follicles and observed a cell autonomous effect that can be quantified by the Tissue OP of the mutant cells versus the WT ones of a same follicle (Fig. 5U-V). Thus, Dys promotes stress fiber orientation in each cell.

### The DAPC and Fat2 have partially redundant functions

We found that the DAPC has only a minor impact on the early phase of stress fiber orientation, with a delayed orientation of actin fibers that is however achieved at stage 8, suggesting alternative mechanisms to orient fibers at these stages (4-8). Similarly, deletion of the intracellular part of Fat2, (*fat2-DeltaICD)* induces a hypomorphic defect with only a slight delay of stress fiber orientation, despite a slower rotation (Aurich and Dahmann, 2016; Barlan et al., 2017; Chen et al., 2017). We therefore wondered what could be the relationship between these two genetic backgrounds. We generated a double mutant background for *fat2/ fat2-DeltaICD* and *Dys* null alleles *(Dys, fat2-DeltaICD / Dys, fat2)* and compared stress fiber alignment for these different genotypes (Fig. 4A-D). Tissue OP analysis shows that single mutants are quite similar and that the double mutants display additive effect (Fig. 4E,I), an additivity that is also observed on follicle elongation (Fig. 4Q). Thus, Dys and Fat2 probably work in parallel for stress fiber orientation once Fat2 has initiated follicle rotation.

**Figure 4:**
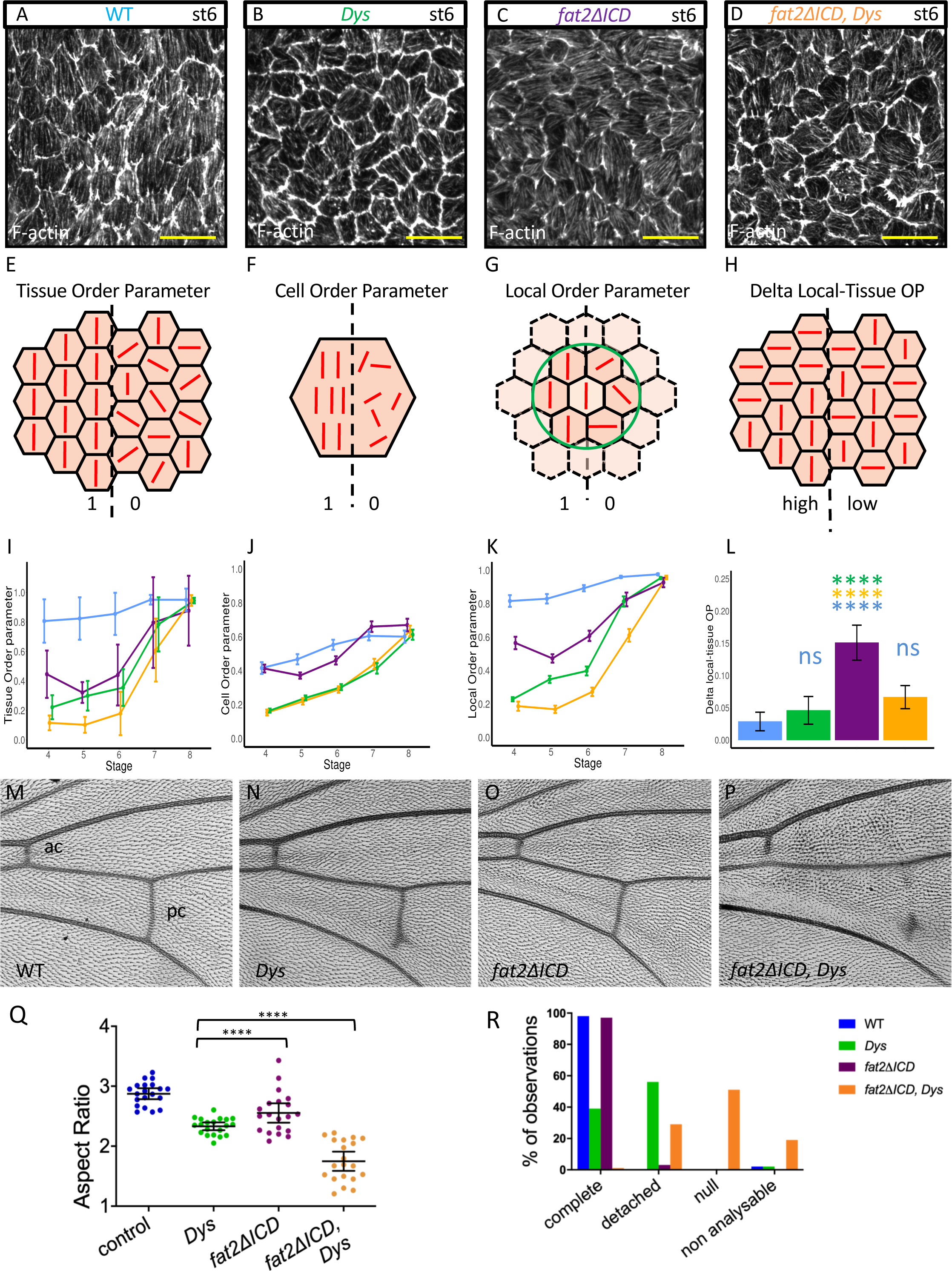
Genetic interaction between *fat2* and *Dys*. A-D) Representative images of stress fibers at stage 6 of A) WT, B) *Dys ^E17^*^/*Exel6184*^ mutant, C) *fat2-DeltaICD* mutant and D) *Dys, fat2-DeltaICD* double mutant. Genotype color code is conserved on all panels. E-H) Schemes explaining the different parameters that are analyzed in I-L) : E,I) Tissue OP (n > 7 follicles for each stage and each genotype): F, J) Cell OP, G,K) Local OP H,L) difference between Local and Tissue OPs (n > 171 cells for each stage and each genotype). P-values between the different genotypes for I,J,K are indicated in supplementary table S3. M-P) Representative images of wing veins of M) WT, N) *Dys ^E17^*^/*Exel6184*^ mutant, O) *fat2-DeltaICD* mutant and P) *Dys, fat2-DeltaICD* double mutant (pc: posterior crossvein, ac: anterior crossvein). Q) Quantification of egg elongation for the indicated genotypes (n>20 eggs for each genotype). R) Quantification of the wing defect observed in the different genotypes on the pc. Non analyzable corresponds to wrinkled or misfolded wings for which crossveins cannot be observed (n=40 wings). For all panels error bars represent s.d.; p *< 0.01, **<0.005, ***<0.001, ****<0.0001. Scalebar 10µm.

**Figure 5:**
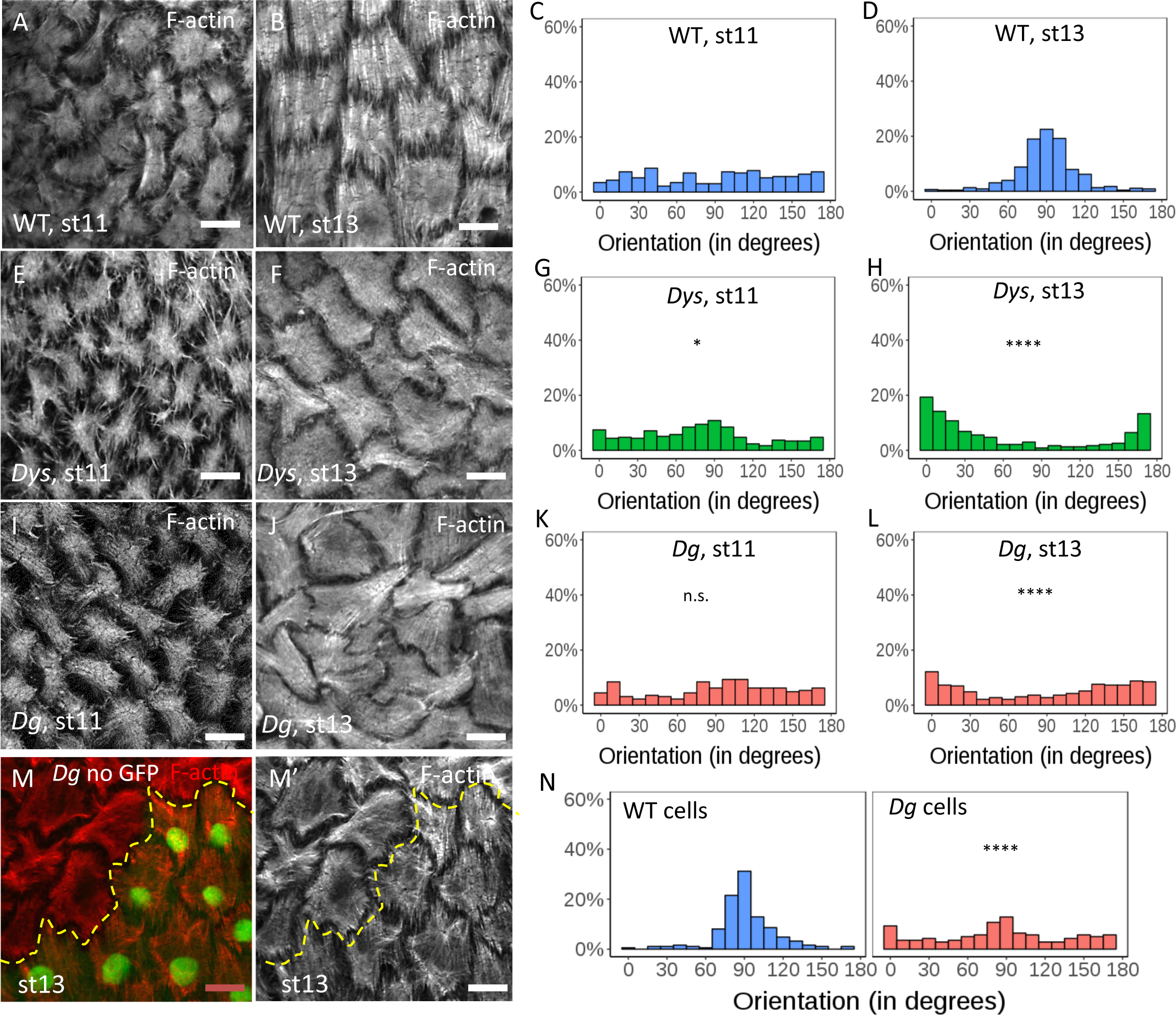
Stress fiber orientation defect in DAPC mutants during late stages. Representative images of basal F-actin in A,B) WT, E,F) *Dys^E17/Exel8164^* and I,J) *Dg^O86^/^O43^* follicles at stage 11 and 13 and C,D,G,H,K,L) Quantification of the corresponding angular distribution (n> 5 follicles). M) Representative image of basal F-actin in *Dg* mutant clone marked by the absence of GFP and N) quantification of the corresponding angular distribution of stress fibers in the mutant cells and the neighbouring WT cells (n> 5 clones). Angular distribution was statistically compared to the WT one of the same stage; p *< 0.01, **<0.005, ***<0.001, ****<0.0001. Scalebar 10µm

To get a better understanding of this additivity we performed a deeper analysis of stress fiber orientation. First, we wondered whether the alignment robustness was the same in each cell and we therefore analyzed the Cell Order Parameter (Cell OP) (Fig. 4F). This clearly shows that the mutants behave in a different manner, with *fat2-DeltaICD* behaving quite similarly to WT whereas *Dys* and *Dys, fat2-DeltaICD* giving an identical weaker Cell OP (Fig. 4J). This effect on Cell OP of *Dys* can be confirmed by its analysis on *Dys* mutant clones (Fig. S4E). Moreover, a close observation of *Dys* single cells reveals more divergent stress fibers than we have observed in *fat2-DeltaICD* cells (Fig. S4C-D). This indicates that Dys promotes stress fiber alignment in each cell, a process that does not seem to involve Fat2. We also calculated a Local Order Parameter (Local OP), determined for each cell on a cluster including this cell and its neighboring cells included in a given radius (Fig. 4G). Globally, for all mutant genotypes that we analyzed, Local OP values are initially higher than the Tissue OP ones and tend to them when increasing the radius (Fig. S4F). Moreover, the additivity of *Dys* and *fat2* can be also observed on this parameter (Fig. 4K and S4F). Finally, we looked at the difference between Local OP and Tissue OP (Fig. 4H). Interestingly, we noticed a stronger Local/Tissue OP difference for *fat2-DeltaICD* than for *Dys* mutants, a difference culminating at stage 6, indicating that stress fiber orientation is less affected at a small scale in *fat2* than in *Dys* mutants (Fig. 4L and S4G). This difference can be also directly observed on images of F-actin orientation (Fig. S4A-B). For *fat2-DeltaICD*, several patches of neighboring cells with a same orientation but changing from a patch to another can be observed. For *Dys*, we did not detected these patches but changes in orientation from one cell to another. Altogether, these data indicate that Dys promotes stress fiber planar polarization at a cellular and local scale whereas Fat2, or at least its intracellular domain, acts at a more global scale.

These results indicate an additive effect of the loss of the DAPC and of this hypomorphic condition for *fat2* and that they are partially redundant to orient stress fibers during early stages. They further suggest that Fat2 and the DAPC are part of a same functional network during follicle elongation, although they probably work in parallel once BM fibrils are deposited.

Intriguingly, *Dys, Dg* and *fat2* mutants show a very specific defect in the wing, with a partial absence of the posterior crossvein, although the cellular basis for this defect is unknown (Christoforou et al., 2008; Viktorinová et al., 2009) (Fig. 4M, N, R). However, this defect is not observed in a *fat2 / fat2-DeltaICD* hypomorphic background (Fig. 4O, R). In the double mutant *Dys, fat2-DeltaICD* flies, the posterior crossvein is usually completely absent or limited to a dot and the anterior crossvein, which is never affected in *Dys* mutant flies, is incomplete (Fig. 4P, R). These defects indicate a strong genetic interaction in the wings as observed in follicles. Thus, these results tend to confirm that Fat2 and the DAPC work in a same network, which might be involved in many contexts.

### The DAPC is required for stress fiber orientation during late stages

Elongation curves of *Dys* and *Dg* mutants indicate that the major elongation defect appears after stage 12 (Fig. 1H). Of note, for unknown reasons, stress fibers lose their planar polarization at stage 10B/11 but regain their proper orientation around stage 12 (Delon and Brown, 2009; Wahlström et al., 2006) (Fig. 5A,B). In WT follicles, this transition can be observed by the change in angular distribution between stage 11 and stage 13 (Fig. 5C,D). F-actin staining of *Dys* and *Dg* null mutant follicles indicates that the stress fibers lose their orientation at stage 11 as in WT, but fail to regain their proper orientation at stage 13 (Fig. 5E,F,I,J). Angular distribution quantification even indicates a tendency of the stress fibers to be aligned parallel to the AP axis at stage 13 (Fig. 5G,H,K,L). Despite their misorientation, these fibers appear normal in structure with, for instance, a correct localization of integrins at their extremities (Fig. S5A-C). Importantly, mitotic recombination to generate mosaic tissues indicates that this function is cell autonomous for *Dg* (Fig. 5M,N), with stress fibers being misaligned only in the mutant cells, as previously observed in *Dys* clones in early stages. The defect is strictly cell autonomous as it is also observed in mutant cells that are adjacent to WT cells. Moreover, WT cells that are adjacent to mutant cells have stress fibers properly oriented (Fig. 5M). Importantly, this cannot be a consequence of a primary effect on BM fibril deposition because follicle rotation has moved the position of the clone with respect to the BM. It reveals, therefore, another function of the DAPC during follicle development. Given the known link of the DAPC with F-actin, it suggests a direct role in stress fiber orientation, sensing and transmitting a cue defining this planar polarization.

### Late DAPC function relies on both early and late activities

Since *Dys* and *Dg* mutants affect BM fibril deposition and stress fiber orientation, we aimed to determine which of these defects is the main contributor to the follicle elongation phenotype. To address this question, we took advantage of the temporality of the DAPC function during oogenesis, being required for BM fibril deposition during early stages and for stress fiber orientation mainly during later stages. Importantly, these three defects, follicle elongation, BM fibril deposition and stress fiber orientation are recapitulated when RNAi against Dg is induced in follicle cells during all stages using tj:Gal4 driver, expressed in follicle cells throughout oogenesis (Fig. 6F-I). We also used Cy2:Gal4 which is only expressed in follicle cells from stage 9 (Fig. S6N). The expression of *Dg* RNAi with this driver knocks it down after the correct BM fibril deposition (Fig. S6A,B), but before the second phase of stress fiber orientation. As expected, the stress fiber orientation is disrupted at stage 13, confirming that this DAPC function is distinct from the one influencing the BM (Fig. S6C,D). Importantly, egg elongation is also affected (Fig. S6E). Thus, BM fibrils are not sufficient to drive late follicle elongation and stress fiber orientation is necessary for it.

**Figure 6:**
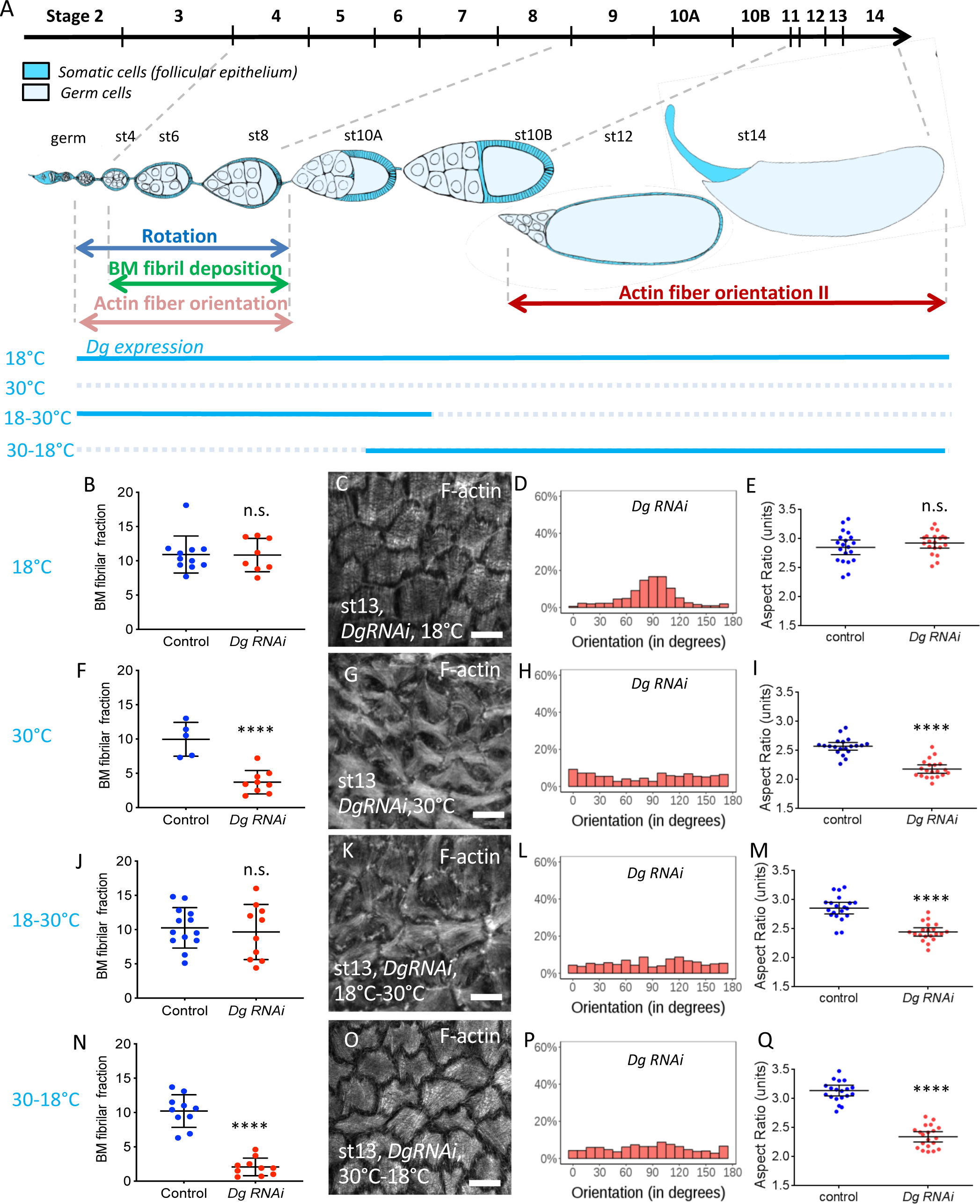
The DAPC is required at two time frames for follicle elongation. A) Scheme of an ovariole with the main events involved in follicle elongation. The above line indicates the time scale of the different developmental stages. The ovariole is oriented anterior to posterior (germ: germarium). Level on endogenous Dg is temporally controlled by the temperature (blue lines). B,F,J,N) Quantification of the fibril fraction of *Dg* RNAi stage 12 follicles expressing ColIV-GFP in the indicated condition compared to control performed in the same temperature condition (n>5 follicles). C,G,K,Q,O) Representative images of basal F-actin of *Dg* RNAi stage 13 follicles in the indicated condition and D,H,L,P) quantification of the corresponding angular distribution (n>5 follicles). E,I,M,Q) Aspect-Ratio quantification of temporally controlled *Dg* RNAi mature compared to control performed in the same temperature condition (n>20 eggs). (For all panels error bars represent s.d.; p *< 0.01, **<0.005, ***<0.001, ****<0.0001)

We also performed the reverse experiment by trying to rescue the late stress fiber orientation defect in a *Dg* mutant using an UAS:Dg. Expression of Dg during the whole oogenesis in the follicle cells, using tj:Gal4 driver in a mutant background, rescues the elongation defect, the BM fibril deposition and stress fiber orientation (Fig. S6F,H,I,J,M). We then tried to rescue *Dg* mutant using the Cy2:Gal4 driver, which clearly allows Dg detection from stage 9 with an antibody failing to detect endogenous level of the protein (Fig S6N. S6O-P). As expected, such follicles show BM defects associated with *Dg* mutation (Fig. S6G,H). However, surprisingly, this genetic combination also fails to rescue stress fiber orientation (Fig. S6K,L). Moreover, egg elongation is only poorly rescued (Fig. S6M). Together these results indicate that late expression of Dg is not sufficient to restore its late functions, and, thus, that these late functions also rely on a previous role of Dg during oogenesis.

To confirm these important observations we set up an alternative method to control the timing of Dg inhibition by RNAi, using Tj:Gal4 and Tub:Gal80^ts^, which is a thermosensitive Gal4 inhibitor allowing a temporal control of its activity. We defined two conditions allowing either Gal4 activity during early stages (switch from 30 to 18°C, “Early *Dg* RNAi”) or late stages (switch from 18 to 30°C, “Late *Dg* RNAi”) (Fig. 6A). Moreover, positive and negative controls were performed at 18°C and 30°C (Fig. 6). For each condition, an internal negative control (driver line) with the same temperature was performed. *Dg* RNAi at 18°C is similar to control (Fig. 6 B-E) whereas at 30°C it recapitulates all the defects observed in *Dg* mutants, as previously mentioned (Fig. 6F-I). The best timing for the temperature switches was adjusted by checking their effect on fibril in late stages, taking into account that these fibrils are formed during early stages. Thus, fibrils are normal in 18-30°C conditions and affected in 30-18°C conditions (Fig. 6J,N). We observed that F-actin orientation at late stages and follicle elongation are affected in both late (Fig. 6K-M) and early (Fig. 6O-Q) *Dg* RNAi conditions, confirming our previous results. Thus, the DAPC is required at two different time windows and its second function is dependent on the first one.

### BM fibril deposition provides a persistent cue for F-actin orientation

Overexpression of *Rab10* increases the BM fibril fraction and enhances follicle elongation, a phenotype opposite to the loss of function of the DAPC (Isabella and Horne-Badovinac, 2016). We therefore performed an epistasis test between these two conditions. Concomitant overexpression of *Rab10* and knock-down of *Dys* in follicle cells using tj:Gal4 increase the amount of BM fibrils to a higher level than the WT control and thus fully compensate for the absence of Dys (Fig. S7A-D, E). This rescue is not due to titration of the Gal4 protein by the second UAS line when we used both *Dys* RNAi and Rab10 UAS constructs as the BM fibril fraction in a UAS: tomato; *Dys* RNAi control is not increased (Fig. S7I). Classical interpretation of such a test would indicate that Rab10 acts downstream (or in parallel) of the DAPC. These experiments were performed at optimal temperature for Gal4-induced expression (30°C). However, in such context *Rab10* overexpression disrupts follicle elongation (Isabella and Horne-Badovinac, 2016), and it is also associated with the abnormal targeting of ColIV containing vesicles to the apical side of the cells (Fig. S7H). Therefore, these conditions did not allow us to determine whether it could also rescue follicle elongation. As a consequence, we repeated these experiments at 25°C and observed an almost complete rescue of BM fibril fraction (Fig. 7A). However, mature egg elongation was still disrupted, as in DAPC loss of function (Fig. 7B). Thus, for elongation, Dys should be placed downstream of Rab10. Taken together, these results are consistent with the proposal that the DAPC has two separate functions, one upstream and one downstream of Rab10. Moreover, it also confirms that, in absence of the DAPC, BM fibrils are not sufficient for elongation.

**Figure 7:**
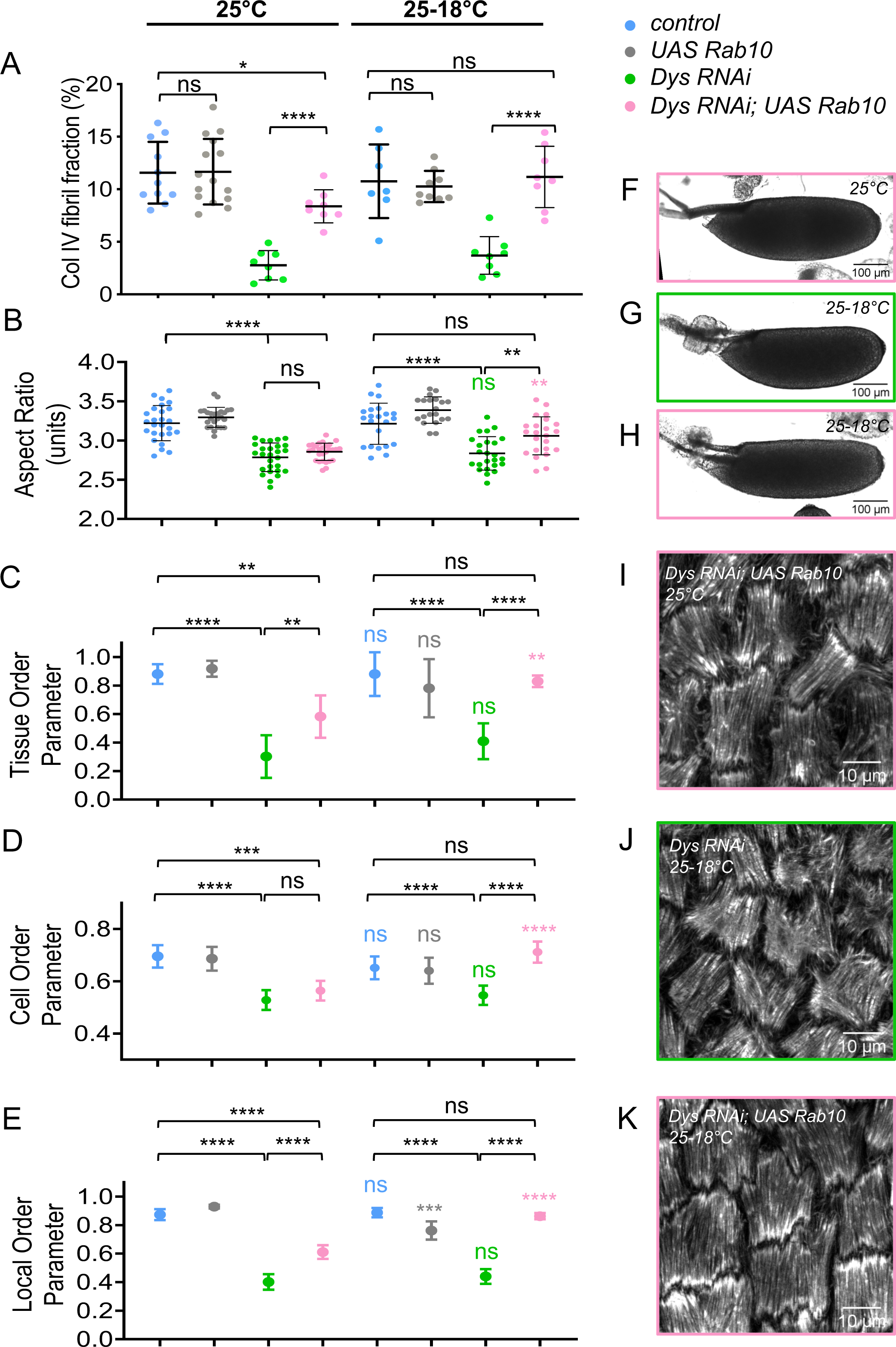
BM fibrils provide a cue for DAPC dependent F-actin orientation. A) Quantification of the BM fibril fraction at stage 8 for the indicated genotypes at 25°C (n>8). B-E) For the indicated genotypes without (25°C) or with (25-18°C) a switch to non-permissive temperature, quantification of B) the Aspect-Ratio of mature eggs C) the Tissue-OP (n > 7 follicles) D) the Cell-OP, and E) the Local-OP at stage 13 (n > 45 cells). F-K) Representative images of F-H) mature eggs and of I-K) stress fibers at stage 13 for the indicated genotypes without (25°C) or with (25-18°C) a switch to non-permissive temperature. For all panels error bars represent s.d.; p *< 0.01, **<0.005, ***<0.001, ****<0.0001. Coloured stars (or ns) correspond to the statistical test for a same genotype between 25°C and 25-18°C conditions.

We also analyzed in this context stress fiber orientation in late stages. Angular distribution for *Dys* RNAi (Fig. S7J), compared to the one of null mutants (Fig. 5H), indicates that it corresponds to hypomorphic conditions. This result prompted us to also analyze the different order parameters as previously done on early stages to get a more precise evaluation of stress fiber orientation and the three were strongly affected in *Dys* RNAi (Fig. 7C-E). When Rab10 is overexpressed at the same time we observed a partial rescue of this defect on the angular distribution, the Tissue OP, the Local OP but not the Cell OP (Fig. S7J, Fig. 7C-E,I). These observations could be explained by the rescue of the early function of the DAPC by Rab10 and the hypomorphic nature of *Dys* RNAi. Nonetheless, this approach offers a unique opportunity to test whether the DAPC’s first function in BM fibril deposition is required for its later function in stress fiber orientation. We repeated the experiments but allowing *Dys* expression during late stages by a temperature switch (25-18°C). Strikingly, the late expression of Dys promotes correct stress fiber orientation on all the parameters that we analyzed only if the BM fibrils have been restored by *Rab10* overexpression (Fig. 7C-E,J,K, Fig. S7J). Moreover, this condition also restores egg elongation (Fig. 7B,F-H). These experiments demonstrate the double function of the DAPC, with the latter in F-actin organization depending on the first one in BM structure. It also shows that the BM fibrils are required for F-actin orientation long after their deposition and that this function explains their contribution to follicle elongation.

## Discussion

In this article, we show that the DAPC contributes to follicle elongation by two distinct functions in which the second relies on the output of the first one. These results are important for understanding the activity of this key complex and the morphogenetic mechanisms involving the ECM.

### Follicle elongation relies on multiple mechanisms

Genetic data already showed that follicle elongation relies on at least two different and successive mechanisms. The first one is controlled by JAK-STAT and involves the follicle cell apical domain, whereas the second one is controlled by Fat2 and involves their basal domain and the BM (Alégot et al., 2018; Aurich and Dahmann, 2016). Between these phases, around stage 7-8, JAK-STAT and Fat2 seem to be integrated in a third mechanism based on a BM stiffness gradient (Crest et al., 2017). Interestingly, a very recent report suggests that this gradient may not directly influence tissue shape but rather do so by modifying the properties of the follicle cells underneath (Chen et al, 2019). Here, we show that the DAPC influences elongation mainly at very late stages, suggesting the existence of a fourth mechanistic elongation phase. Of note, elongation at these late stages is also defective in *fat2* mutants (Alégot et al., 2018; Aurich and Dahmann, 2016). This is consistent with the fact that rotation is required for polarized BM fibril deposition, and that this deposition depends on and is required for DAPC function. The existence of multiple and interconnected mechanisms to induce elongation, a process that initially appeared to be very simple, highlights the true complexity of morphogenesis, and the necessity to explore it in simple models.

### BM fibrils act as a planar polarity cue and F-actin as a molecular corset

Fat2 is clearly part of the upstream signal governing the basal planar polarization. However, how this polarization leads to tissue elongation is still debated. It has been proposed that elongation relies on a molecular corset that could be formed, non-exclusively, by BM fibrils or F-actin stress fibers (Gutzeit, 1990; Gutzeit et al., 1991; Haigo and Bilder, 2011; Frydman and Spradling, 2001; Bateman et al., 2001; Viktorinová et al., 2009; Cetera et al., 2014). The initial observation that rotation is required for both elongation and BM fibrils favored a direct mechanical role for these structures (Haigo and Bilder, 2011). Recent data showing that BM fibrils are stiffer than the surrounding ECM supports this view (Chlasta et al., 2017). Moreover, increasing the fibril number and size can lead to hyper-elongation (Isabella and Horne-Badovinac, 2016). Finally, addition of collagenase induces follicle rounding, at least at some stages, and genetic manipulation of the ECM protein levels also influences elongation (Haigo and Bilder, 2011; Isabella and Horne-Badovinac, 2015; Chlasta et al., 2017). However, these experiments did not discriminate between the function of the fibril fraction and a general BM effect. Moreover, they do not allow to discriminate whether their impact on elongation is direct and mechanic or indirect, by a specific response of the epithelial cells. Fat2 and rotation are also required for the proper orientation of the stress fibers. The F-actin molecular corset is dynamic with follicle cells undergoing basal pulsations, and perturbation of both these oscillations and of the stress fiber structure affect elongation (Bateman et al., 2001; He et al., 2010). In the DAPC mutants we observed a faint but significant elongation defect during mid-oogenesis and a stronger one after stage 12. These defects are clearly correlated with the stress fiber orientation defects observed in the same mutants, both temporally and in terms of intensity. Moreover, rescuing the BM fibrils in a *Dys* loss of function by *Rab10* overexpression does not rescue elongation indicating that stress fiber orientation is instrumental.

Thus, if the role of the BM fibrils as a direct mechanical corset appears limited, what is their function? One possibility could have been that they promote rotation, acting in a positive feed-back and explaining speed increase over time. However, the rotation reaches the same speed in WT and DAPC mutants, excluding this possibility. Similarly, increasing the fibril fraction also has no effect on rotation speed (Isabella and Horne-Badovinac, 2016).

Our results strongly argue that BM fibrils act as a cue for the orientation of stress fibers, which then generate the mechanical strain for elongation. This appears clear in late stages where the function of the DAPC for stress fiber orientation is dependent on the previous BM fibril deposition. Whereas it is unknown why the cells lose their orientation from stages 10 to 12, these stages are associated with a strong remodeling of the follicular epithelium. The BM fibrils provide the long-term memory of the initial pcp of the tissue, allowing stress fiber reorientation (Figure 8). Such mechanism appears as a very efficient way to memorize positional cues, and could represent a general BM function in many developmental processes.

**Figure 8:**
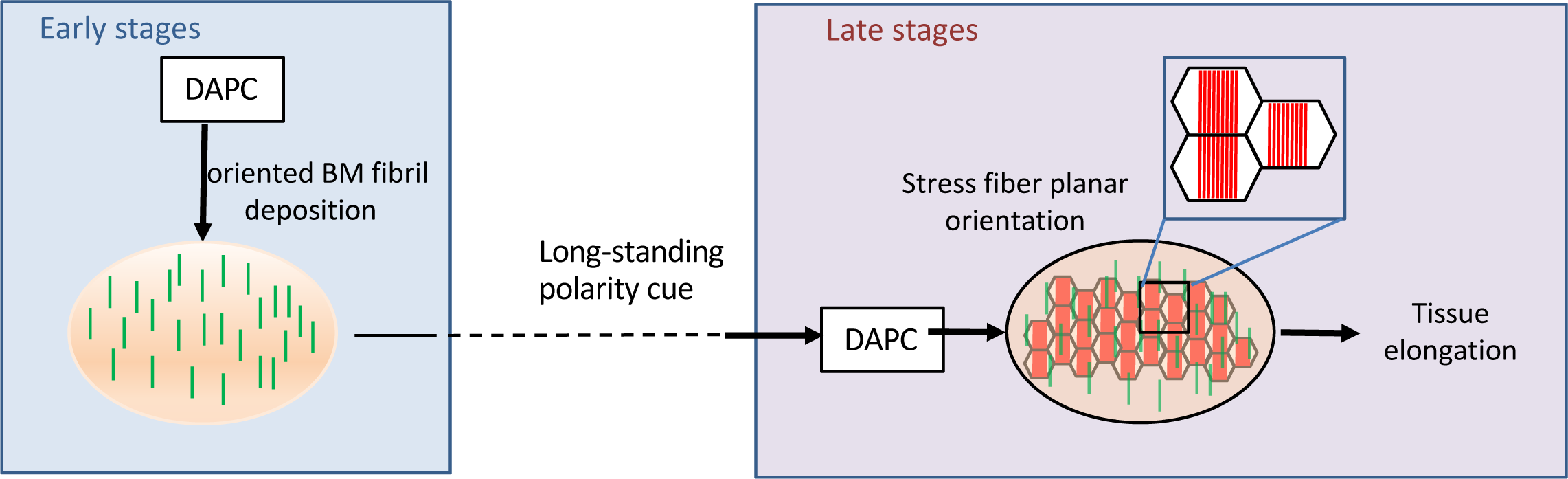
Dual function of the DAPC during elongation revealing that BM fibrils work as a persistent planar polarity cue. During the early stages (4-8) the DAPC participates to the formation of oriented BM fibrils perpendicular to the follicle elongation axis. These fibrils are maintained during follicle development including stages with intense morphogenetic processes unrelated to elongation (stages 9-11). At late stages, cells use the persistent BM fibrils as a signal to guide the planar orientation of the stress fibers. This event depends on a distinct second function of the DAPC. The stress fibers are the main mechanical effector of follicle elongation.

How BM fibrils contribute to stress fiber orientation remains to be elucidated. Nevertheless, we can hypothesize that it involves their higher stiffness or a higher density in binding sites for ECM receptors. Our data indicates that the DAPC might be such a receptor. This hypothesis is supported by our findings that the DAPC acts in a cell autonomous manner and promotes stress fiber orientation in each cell and rather locally than at a tissue scale. Of note, this kind of pcp is unusual because it does not require signal transmission between neighboring cells, usually through heterophilic protein-protein interactions. However, an interesting parallel with the pcp established by Fat/Dachsous and the core pcp pathways could be drawn. In many tissues these two pathways work in parallel and their cooperation is required for proper pcp establishment at the apical domain of epithelial cells (Hale and Strutt, 2015). The genetic interaction that we revealed between Fat2 and the DAPC suggests that robust basal pcp establishment also requires the integration of two independent mechanisms. Importantly this genetic interaction applies to at least two tissues suggesting a general functional link between them.

### The DAPC acts as an organizer of the basal domain of epithelial cells

We found that the DAPC impacts the two key actors at the basal domain of the follicle cells : the BM and the stress fibers linked to the BM. All the defects of *Dg* null mutants were also observed in *Dys* mutants, demonstrating a developmental and morphogenetic role for this gene. In vertebrates, at least in some tissues, Dg presence seems essential for BM assembly (Henry and Campbell, 1998; Satz et al., 2010; Bello et al., 2008; Buisson et al., 2014). However, BM formation on the follicular epithelium does not require Dg, suggesting the existence of alternative platforms for its general assembly. Our genetic data suggest that Rab10 is epistatic to Dg for BM fibril deposition. The usual interpretation of such a result would be that Dg is involved in the targeting of ECM secretion upstream of Rab10 rather than in ECM assembly in the extracellular space. In *C. elegans*, Dg acts as a diffusion barrier to define a precise subcellular domain for ECM remodeling (Naegeli et al., 2017). One could imagine that the DAPC has a similar function in follicle cells, by defining the position where the Rab10 secretory route is targeted. However, in DAPC loss of function, some ECM is still secreted between cells, suggesting that the lateral Rab10 route is not affected. Moreover, ECM proteins do not abnormally accumulate between cells in such mutants, suggesting that they are able to leave this localization but without forming fibrils. Therefore, the functional interplay between Rab10 and the DAPC is still unclear.

As mentioned before, Dg has often been proposed to act as a scaffold to promote BM assembly in mice. Deletion of the Dg intracellular domain is only sub-lethal in mice, whereas complete loss of this protein is lethal very early during development, indicating that the abolishment of Dg’s interaction with Dys affects only partially its function (Henry and Campbell, 1998; Satz et al., 2009). In these mice, laminin assembly can still be observed, for instance in the brain and retina (Satz et al., 2010; Clements et al., 2017). Similar results were also obtained in cultured mammary epithelial cells (Weir et al., 2006). Thus, despite the existence of Dys paralogs that could mask some effects on ECM and the fact that we observed the same ECM alteration in *Dg* or *Dys* mutant fly follicles, not all the Dg functions related to ECM assembly or secretion involve Dys. It is possible that Dys is required when Dg needs a very specific subcellular targeting for its function, whereas a more general role in ECM assembly would be Dys-independent. Our results suggest that some specific Dys effects on ECM could have been underestimated and this could help to explain the impact of its loss of function on tissue integrity maintenance. For instance, as it has been reported that Dg influences ECM organization in fly embryonic muscles, it would be interesting to determine whether this also involves Dys (Yatsenko and Shcherbata, 2014).

The DAPC is involved in the planar polarization of the basal stress fibers and its ability to read ECM structure to orchestrate integrin-dependent adhesion could play a role in many developmental and physiological contexts. The link between ECM and F-actin provided by this complex is likely required for this function, although it remains to be formally demonstrated. BM fibrils could provide local and oriented higher density of binding sites for Dg, and the alignment could then be transmitted to the actin cytoskeleton. Alternatively, DAPC function could rely on sensing the mechanical ECM properties. The hypothesis that the DAPC could act as a mechanosensor is a long standing proposal, partly due to the presence of spectrin repeats in Dys (Brown and Lucy, 1993; Garbincius and Michele, 2015). The basal domain of the follicle cells may offer an amenable model to combine genetics and cell biology approaches to decipher such function.

Altogether, this work provides important insights on BM role during morphogenesis, by acting as a static pcp cue retaining spatial information while cells are highly dynamic. It also reveals important functions of the DAPC, including Dys, that may be broadly involved during animal development and physiology.

## Methods

### Genetics

The detailed genotypes, temperature and heat-shock conditions are given in supplementary table 1.

### Live imaging and rotation speed analysis

For live imaging, ovaries were dissected, as described previously (Alégot *et al*, 2018), and stained with the membrane dye FM4-64. Samples were cultured for less than 1 hour before imaging. Damaged follicles (identified on the basis of abnormal FM4-64 incorporation) were not analyzed. Rotation speed analysis was performed using a Fiji Manual Tracking plug-in. For each follicle, speed was measured at three independent positions during 30min.

### Dissection and immunostaining

Resources and reagents are listed in supplementary table 2.

Dissection and immunostaining were performed as described previously (Vachias et al., 2014) with the following changes : ovaries were dissected in supplemented Schneider medium, and ovarioles were separated and the muscle sheath was removed before fixation to obtain undistorted follicles. Stage was determined using unambiguous reference criteria that are follicle shape-independent (Spradling et al., 1993). For quantification of stress fiber orientation, ovaries were fixed with 8% paraformaldehyde (PFA) containing phalloidin-TRITC or phalloidin Atto-488. They were then permeabilized with PBS-Triton at 0.02% and incubated again with phalloidin-TRITC (or Atto-488). Images were taken using a Leica SP8 or a Zeiss LSM800 Airyscan confocal microscope excepted for stage 12 BM images that were acquired on an inverted Zeiss Spinning Disc (Yokogawa CSU-22). Immunostaining of pericellular ECM was performed as described by Isabella and Horne-Badovinac, 2016, using antibodies against GFP (ColIV-GFP-expressing flies) or against LanA (a gift from Volk’s laboratory).

### Aspect Ratio determination

The length of the long and short axes of each follicle were directly measured by transmitted light microscopy and used to determine the aspect ratio.

### BM fraction, pericellular matrix and F-actin orientation quantification

BM fibril fraction was determined using a homemade Fiji macro, using the principles developed by Isabella and Horne-Badovinac, 2016. Threshold for fibril fraction determination was manually adjusted on each control experiment to obtain a proper fibril detection, due to slight variation of the fluorescence depending on the temperature, the developmental stage and the used microscope. used. The same threshold was then assigned for all the genotypes of the same experiment. The pericellular matrix was quantified using the Fiji tool to measure the “meangray value” of a 1.32µm wide line drawn on the cell walls (about 100 lines for each genotype – 20 cells).

Stress fiber orientation and Tissue OP analysis were performed as described in Cetera et al., 2014 after manual segmentation. Analyses were performed on 40 and 96 μm^2^ images for the early and late stages, respectively. The Cell OP was determined in the same way, by analyzing the alignment robustness at the cellular scale. For Local OP, the average cell radius of each image was determined. Then, we analyzed the cells with their barycenter being in 3.5 radiuses from the considered cell, this value fitting with an average of 6 neighbors. This value included on average six neighboring cells. Clusters with less than four cells were excluded. Then, the radius was progressively increased from 3.5 to 25. At 25, all the cells of the image were included, and thus the Local OP value corresponded to the Tissue OP value. Therefore, this approach was used to observe the tendency from Local OP to Tissue OP and to calculate for each cell the difference between them.

### Statistical analysis

For all experiments, the minimal sample size is indicated in the figure legends. Results were obtained from at least two independent experiments, and for each experiment multiple females were dissected. Randomization or blinding was not performed. The normality of the samples was calculated using the D’Agostino & Pearson normality test. The unpaired t-test was used to compare samples with normal distribution, and the unpaired Mann-Whitney test for samples without normal distribution.

Concerning the stress fiber angular distribution, the circular mean orientation of the fibers was computed for each cell and then used for the Rao’s test of homogeneity. For each stage, a pairwise comparison of genotypes was performed to determine whether the polar vectors or the dispersions were equal. These tests were performed in R using the “circular” package.

## Acknowledgements

We thank R. Ray, H. Ruohola-Baker and T. Volk for fly stocks or reagents. This work was supported by the Association Française contre les Myopathies (AFM) (Trampoline grant 17683 and MyoNeurAlp network). This research was also financed by the French government IDEX-ISITE initiative 16-IDEX-0001 (CAP 20-25) and by a grant from the American Cancer Society (RSG-14-176) to S.H-B. FCC was supported by the Fondation pour la Recherche Médicale (FRM, FDT20170437189), and AI by NIH T32 HD055164 and an NSF Graduate Research Fellowship. We also thank the CLIC facility (Clermont Imagerie Confocale), and team members for comments on the manuscript.

## Supplemental data

**Supplemental figure 2:**
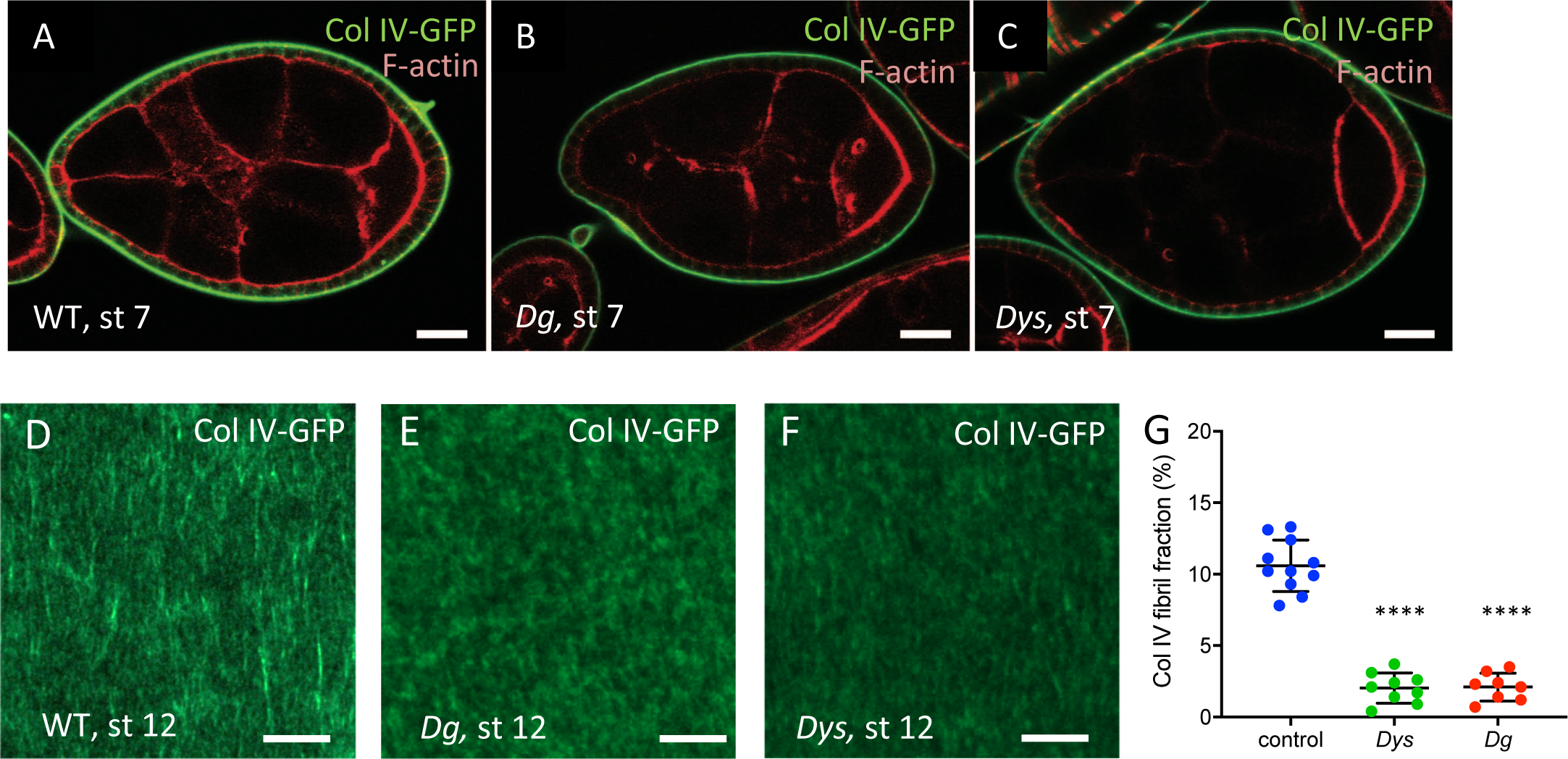
A-C) Optical cross-section of stage 7 WT, *Dys ^E17^*^/*Exel6184*^ and *Dg ^O46/O83^* null mutant follicles that express ColIV-GFP and stained for F-actin. Scale bar ? D-F) Basal view of the ECM of stage 12 WT, *Dys ^E17^*^/*Exel6184*^ and *Dg ^O46/O83^* null mutant follicles that express ColIV-GFP. G) Quantification of the fibril fraction in stage 12 WT, *Dys ^E17^*^/*Exel6184*^ and *Dg ^O46/O83^* null mutant follicles. Scalebar 10µm. For all panels error bars represent s.d.; p *< 0.01, **<0.005, ***<0.001, ****<0.0001.

**Supplemental figure 4:**
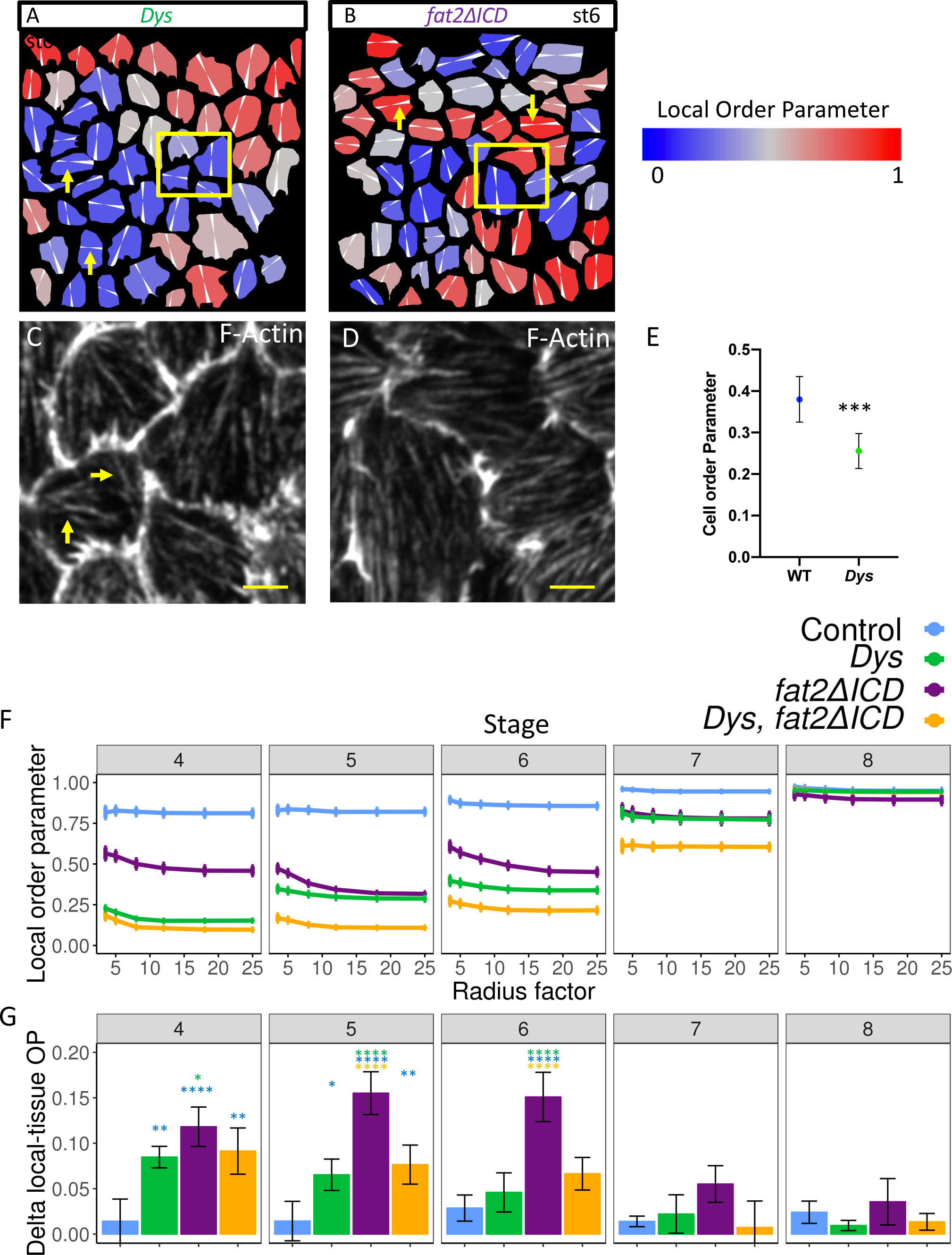
A-B) Color coding of the Local OP by cell on representative images (B and C of Fig. 4) of A) *Dys ^E17^*^/*Exel6184*^ and B) f*at2deltaICD* mutants. Main orientation of the stress fibers is indicated in white. Note that cells with a wrong orientation (yellow arrows) can have a high Local OP in *fat2deltaICD* mutant (but not in *Dys* mutant) indicating that they are surrounded by cells with a similar orientation. C-D) Zoom-in corresponding to the yellow squares in A and B. Yellow arrows highlight divergent stress fibers in a same cell in *Dys* mutant. E) Cell Order Parameter for *Dys* mutant cells and WT neighbouring cells at stage 5 (n >138 cells). F) Representation for stages 4 to 8 of the Local OP as a function of the distance (measured in cell radius), showing the tendency from Local OP to Tissue OP. G) Difference between Local OP (minimal value for radius in E) and Tissue OP (maximal value for radius in E) (n > 171 cells for each stage and each genotype). For all panels error bars represent s.d.; p *< 0.01, **<0.005, ***<0.001, ****<0.0001. Scale bar 2µm

**Supplemental figure 5:**
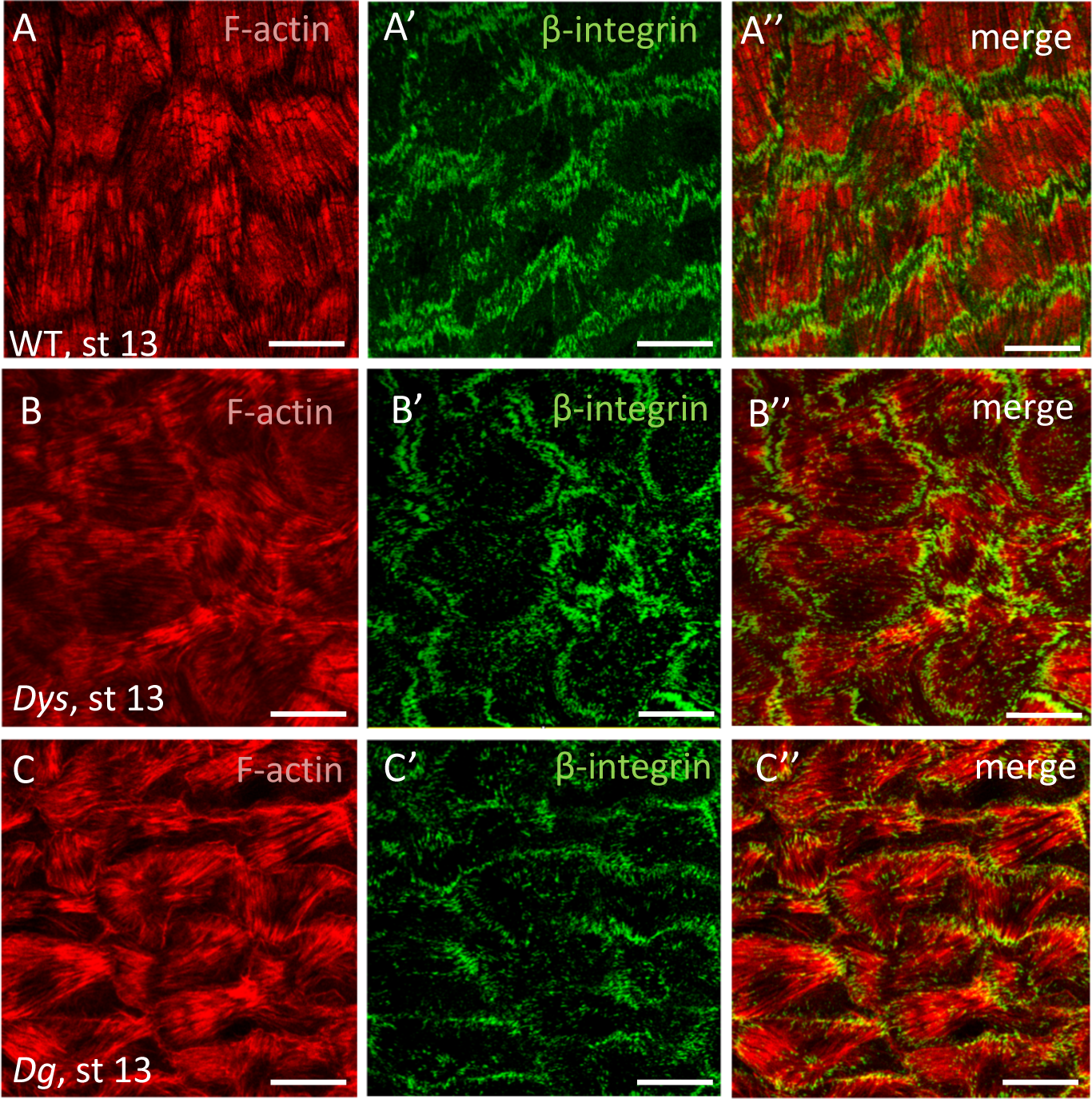
F-actin and Beta-integrin staining in A) WT, B) *Dys ^E17^*^/*Exel6184*^ and C) *Dg ^O46/O83^* mutant follicles at stage 13. Scale bar 10µm

**Supplemental Figure 6:**
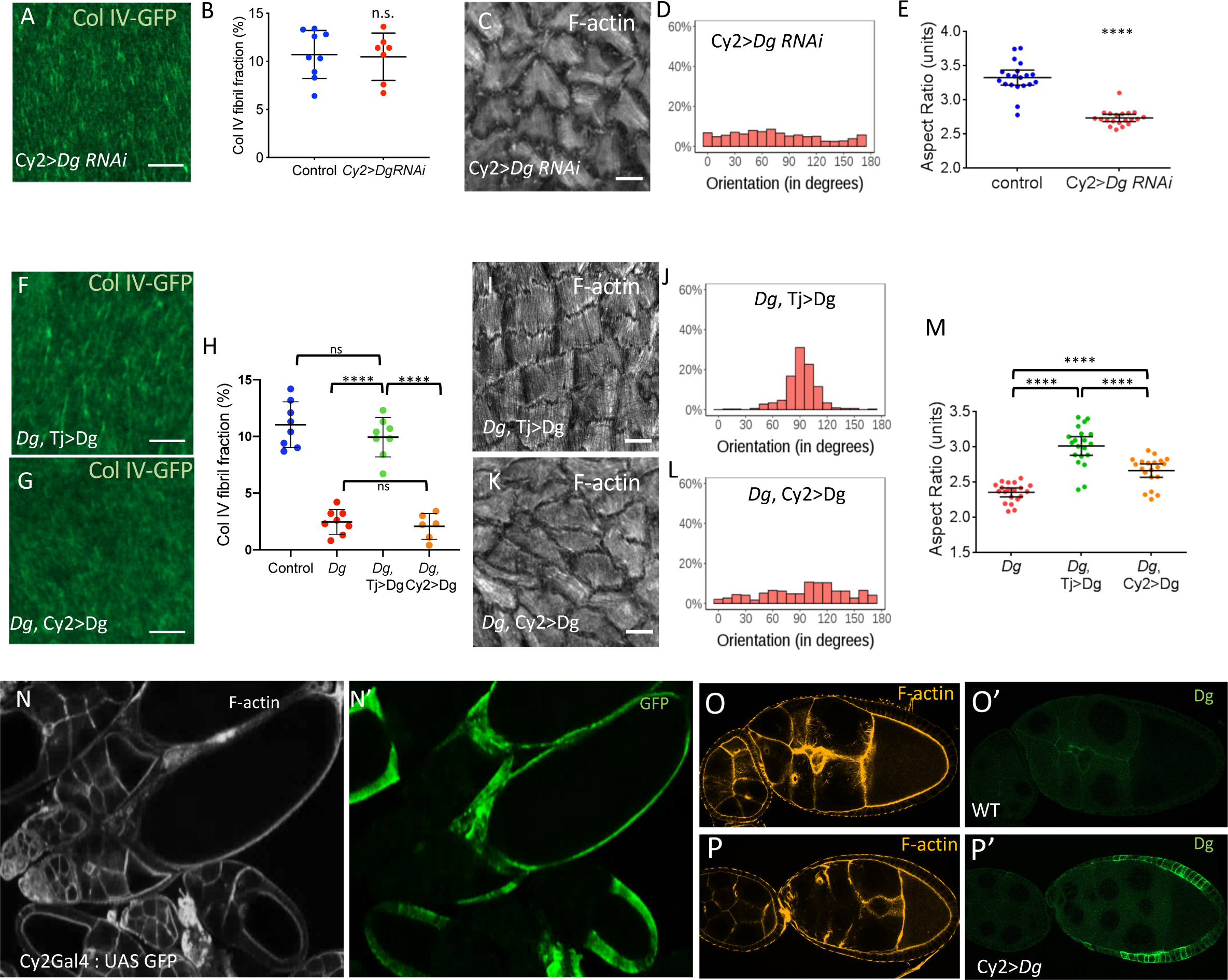
A,F,G) Basal view of the ECM at stage 12 of A) Dg RNAi driven with Cy2:Gal4, F) *Dg ^O46/O83^* mutant expressing UAS:Dg driven with tj:Gal4, G) Dg mutant expressing UAS:Dg driven with Cy2:Gal4 B,H) Quantification of the BM fibril fraction for the indicated genotypes at stages 12. C,I,K) Representative images of basal F-actin at stage 13 of C) Dg RNAi driven with Cy2:Gal4, I) Dg mutant expressing UAS:Dg RNAi driven with tj:gal4, K) *Dg* mutant expressing UAS:Dg RNAi driven with Cy2:Gal4 D,J, L) Quantification of stress fiber angular distribution of the same genotypes than in C,J,K). E, M) Aspect-Ratio quantification of mature eggs of the indicated genotypes. N) Expression profile of Cy2:Gal4. GFP can be detected from stage 9. O-P) Dg immunostaining in WT and Cy2:Gal4 UAS:Dg flies. Endogenous Dg is not detected but induced expression is clearly detectable at stage 9. (For all panels error bars represent s.d.; p *< 0.01, **<0.005, ***<0.001, ****<0.0001) Scale bar 10µm.

**Supplemental figure 7:**
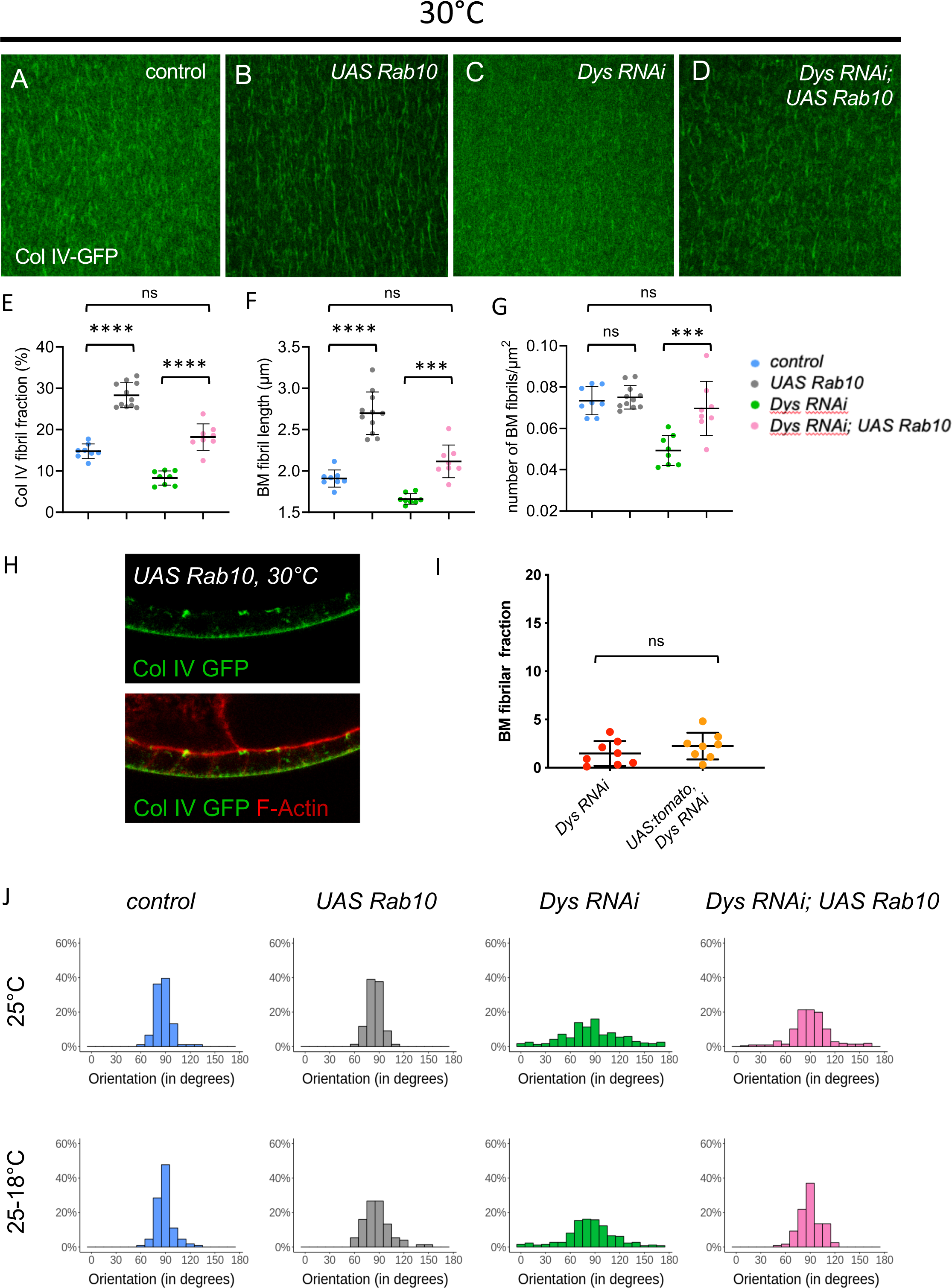
A-D) Basal view of the ECM at stage 8 of A) WT, B) tj:Gal4>UAS:Rab10, C) tj:Gal>UAS Dys RNAi, D) tj:Gal4> UAS Dys RNAi, UAS Rab10. E-G) Quantification of E) BM fibril fraction F) fibril length and G) fibril numbers per µm^2^ at stage 8 for the indicated genotypes at 30°C (n>8). (error bars represent s.d.; p *< 0.01, **<0.005, ***<0.001, ****<0.0001) H) Section view of the follicular epithelium expressing ColIV-GFP and overexpressing Rab10 at 30°C. I) Quantification of fibril fraction at stage 12 for tj>*Dys* RNAi and tj>UAS:tomato; *Dys* RNAi J) Angular distribution for the indicated genotypes without (25°C) or with (25-18°C) a switch to non-permissive temperature.

**Supplementary table 1:**
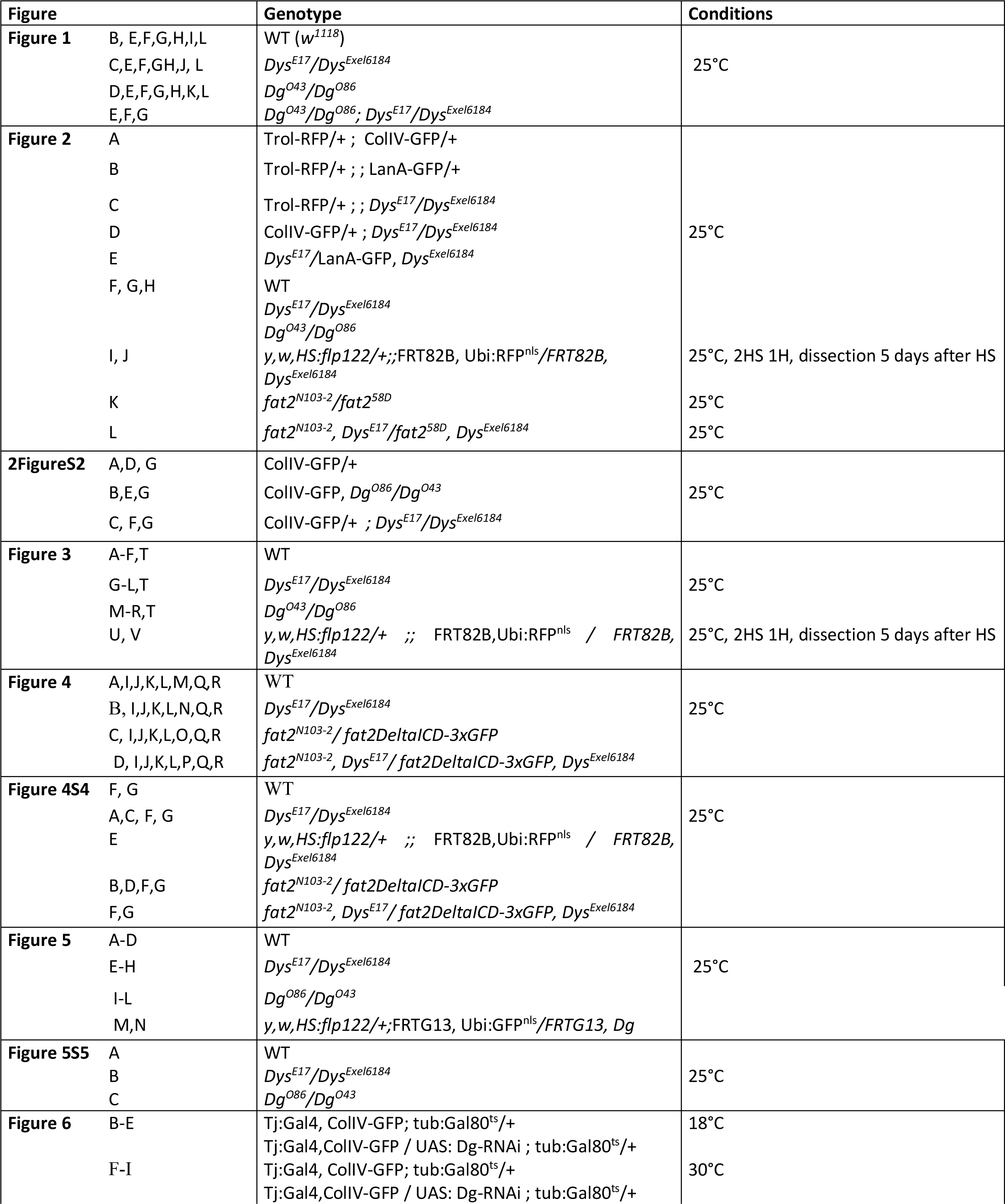
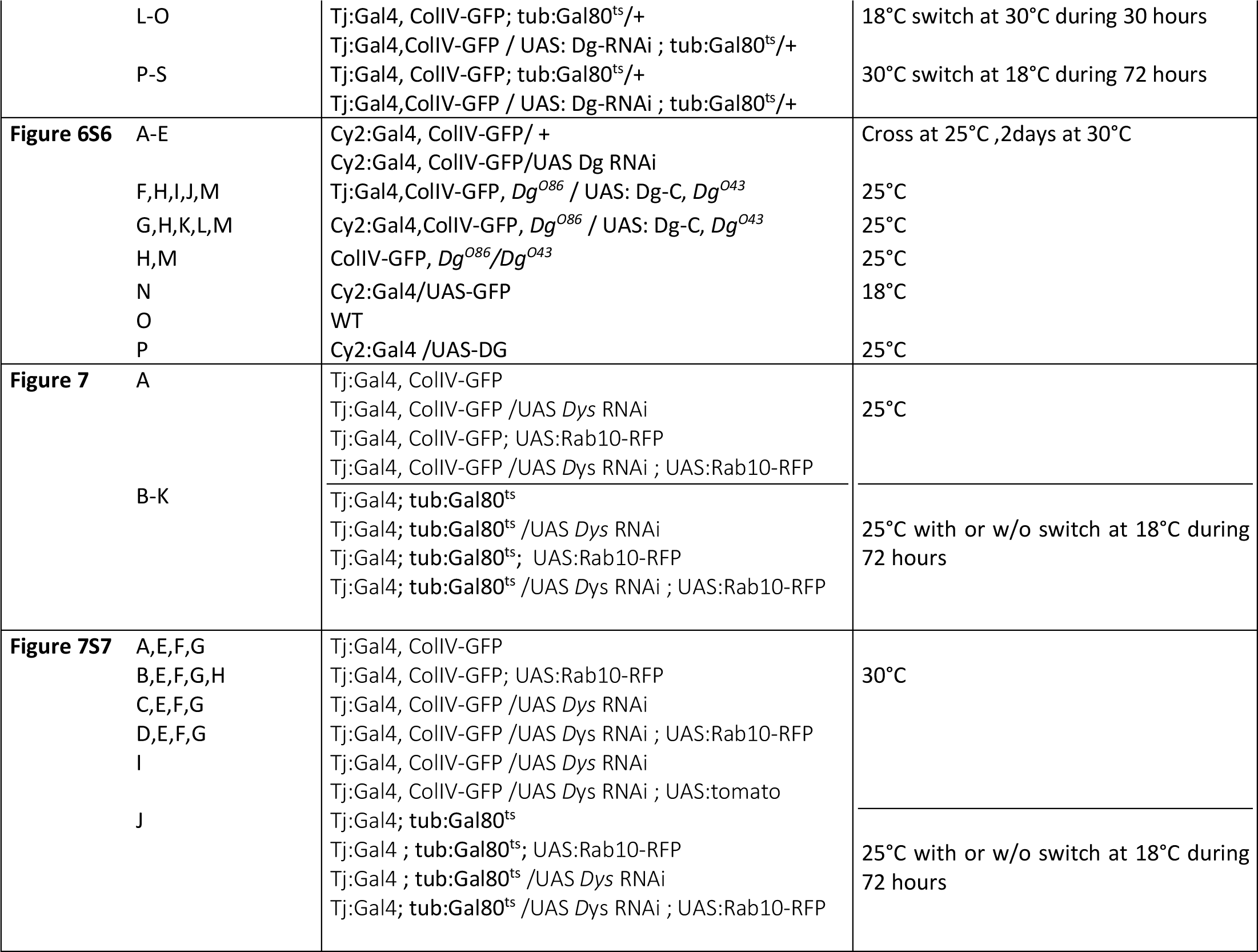
Genotypes and specific conditions.

**Supplementary table 2:**
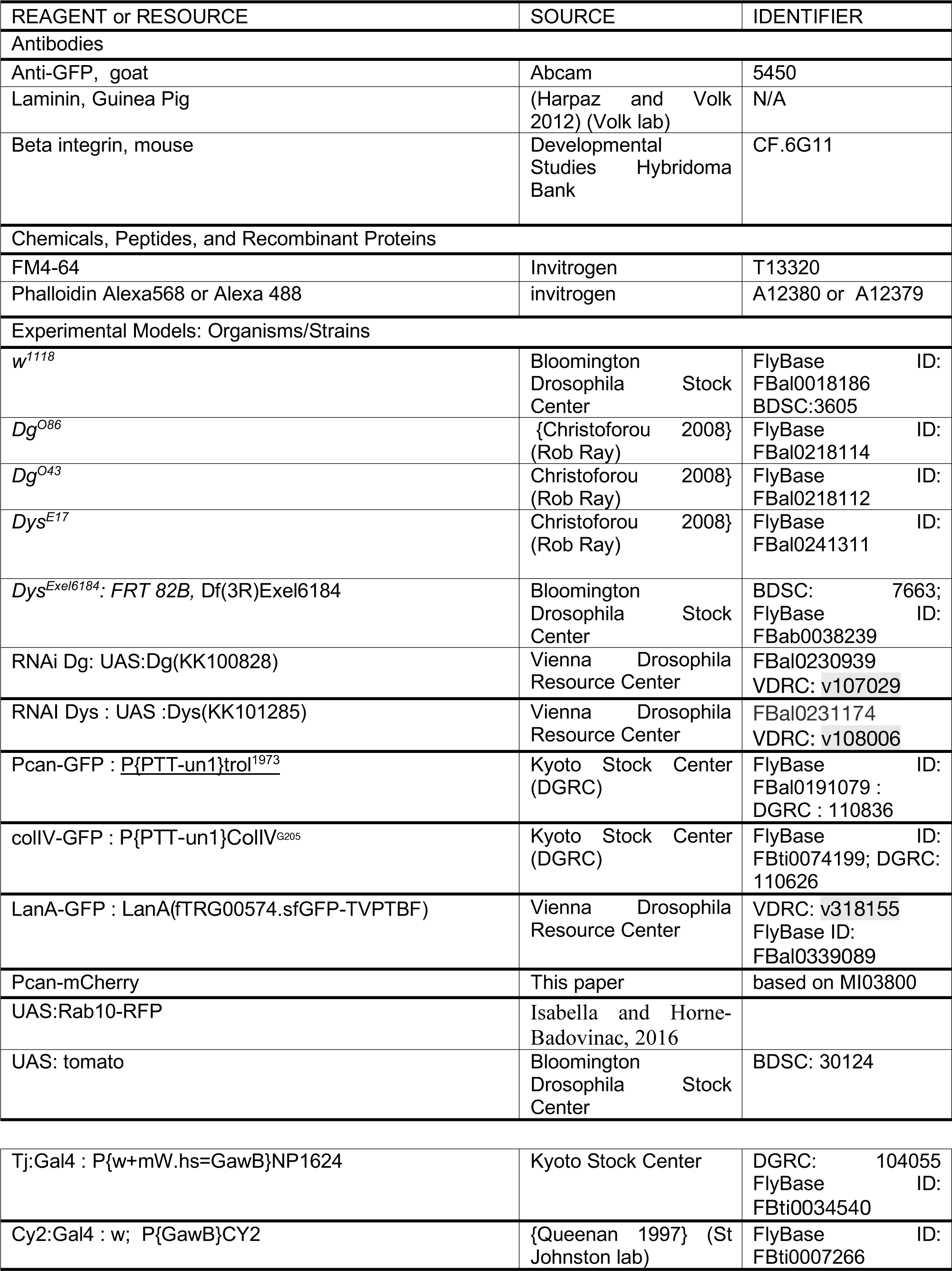
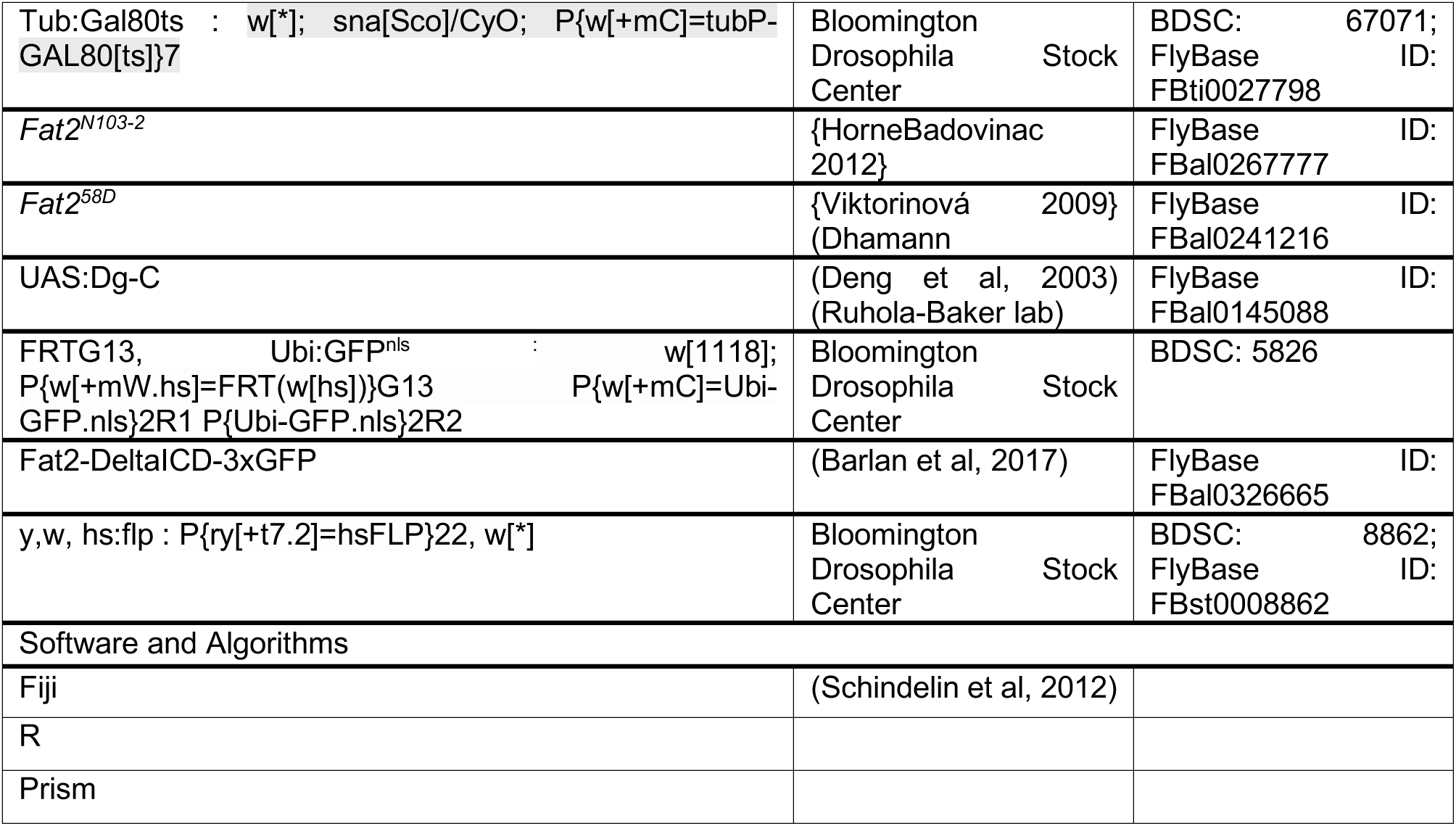
RESOURCES and REAGENT TABLE.

**Supplementary table 3:** Results of the statistical analysis for Fig 7I-L.

| test genotype 1- genotype 2 | LOP<br>p value | COP<br>p value | TOP<br>p value | Delta LOP -TOP<br>p value |
| --- | --- | --- | --- | --- |
| Stage 4: |  |  |  |  |
| WT-Fat2ΔICD | <.0001 | 0.9990 | 0.0008 | <.0001 |
| WT-Dys,Fat2ΔICD | <.0001 | <.0001 | <.0001 | 0.0038 |
| WT-Dys | <.0001 | <.0001 | <.0001 | 0.0023 |
| Fat2ΔICD-Dys,Fat2ΔICD | <.0001 | <.0001 | <.0001 | 0.3807 |
| Fat2ΔICD-Dys | <.0001 | <.0001 | 0.0055 | 0.0452 |
| Dys,Fat2ΔICD-Dys | 0.0613 | 0.8408 | 0.4865 | 0.9629 |
| Stage 5: |  |  |  |  |
| WT-Fat2ΔICD | <.0001 | <.0001 | <.0001 | <.0001 |
| WT-Dys,Fat2ΔICD | <.0001 | <.0001 | <.0001 | 0.0059 |
| WT-Dys | <.0001 | <.0001 | <.0001 | 0.0166 |
| Fat2ΔICD-Dys,Fat2ΔICD | <.0001 | <.0001 | 0.0250 | <.0001 |
| Fat2ΔICD-Dys | <.0001 | <.0001 | 0.9973 | <.0001 |
| Dys,Fat2ΔICD-Dys | <.0001 | 0.3865 | 0.0413 | 0.8432 |
| Stage 6: |  |  |  |  |
| WT-Fat2ΔICD | <.0001 | <.0001 | 0.0004 | <.0001 |
| WT-Dys,Fat2ΔICD | <.0001 | <.0001 | <.0001 | 0.1535 |
| WT-Dys | <.0001 | <.0001 | <.0001 | 0.7643 |
| Fat2ΔICD-Dys,Fat2ΔICD | <.0001 | <.0001 | 0.0209 | <.0001 |
| Fat2ΔICD-Dys | <.0001 | <.0001 | 0.8242 | <.0001 |
| Dys,Fat2ΔICD-Dys | <.0001 | 0.6676 | 0.1256 | 0.3830 |
| Stage 7: |  |  |  |  |
| WT-Fat2ΔICD | <.0001 | 0.0549 | 0.5779 | 0.2422 |
| WT-Dys,Fat2ΔICD | <.0001 | <.0001 | 0.0079 | 0.9879 |
| WT-Dys | <.0001 | <.0001 | 0.5052 | 0.9830 |
| Fat2ΔICD-Dys,Fat2ΔICD | <.0001 | <.0001 | 0.3924 | 0.0844 |
| Fat2ΔICD-Dys | 0.9693 | <.0001 | 1.0000 | 0.4280 |
| Dys,Fat2ΔICD-Dys | <.0001 | 0.5118 | 0.4610 | 0.8844 |
| Stage 8: |  |  |  |  |
| WT-Fat2ΔICD | 0.4559 | 0.0574 | 0.9600 | 0.9742 |
| WT-Dys,Fat2ΔICD | 0.9388 | 0.5843 | 1.0000 | 0.9784 |
| WT-Dys | 0.9164 | 0.9753 | 1.0000 | 0.9538 |
| Fat2ΔICD-Dys,Fat2ΔICD | 0.7539 | 0.4490 | 0.9685 | 0.8093 |
| Fat2ΔICD-Dys | 0.8422 | 0.0979 | 0.9620 | 0.7549 |
| Dys,Fat2ΔICD-Dys | 0.9995 | 0.7944 | 1.0000 | 0.9986 |

